# The acetylation of the histone-like protein HBsu at specific sites alters gene expression during sporulation in *Bacillus subtilis*

**DOI:** 10.1101/2025.05.07.652692

**Authors:** Liya Popova, Hritisha Pandey, Olivia R. Schreiber, Charalampos Papachristou, Valerie J. Carabetta

## Abstract

Sporulation is an adaptive response to starvation in bacteria that consists of a series of developmental changes in cellular morphology and physiology, leading to the formation of a highly resistant endospore. In *Bacillus subtilis*, there is an intricate developmental program which involves the precise coordination of gene expression and ongoing morphological changes to yield the mature spore. The histone-like protein HBsu is involved in proper spore packaging and compaction of the chromosomal DNA. Previously, we found that the acetylation of different lysine residues on HBsu impairs sporulation frequency and spore resistance properties. One mechanism by which HBsu influences the process of sporulation could be through the regulation of gene expression. To test this idea, we performed RT-qPCR to analyze gene expression throughout the sporulation process in wildtype and seven acetylation-mimicking (glutamine substitutions) mutant strains. Acetylation of HBsu at K41 increased the expression of key early and late sporulation genes, especially during the later stages. For example, overexpression of σ^F^ and σ^G^ drive expression of their regulon members at inappropriate times. These findings suggest that K41 acetylation activates gene expression and might represent an “on-off” switch for important regulatory factors as cells transition from early to late phases. The gene expression profiles of *hbsK3Q*, *hbsK37Q, hbsK75Q, hbsK80Q,* and *hbsK86Q* mutants were largely unchanged, but did have significant reductions of key late sporulation proteins, which could explain the observed defects in spore resistance properties. We propose that acetylation of HBsu at specific sites directly regulates gene expression during sporulation and this is required for proper timing and coordination.

## Introduction

Sporulation is an adaptive response to starvation in bacteria that consists of a series of developmental changes in cellular morphology and physiology, leading to the packaging of one cell into a highly resistant spore (1, 2, 3). The process can be divided into seven stages based on morphology (4). It is important to note that the sporulation stages are usually described as discrete morphological events, whereas the process is actually continuous and several steps occur concomitantly (5). No morphological changes at stage 0 of sporulation occur within the cell, but the decision to sporulate has already been made with high levels of the activated form of the master transcriptional regulator Spo0A (6). Activation of Spo0A occurs through a multistep phosphorelay (7, 8) and variation of phosphorylated Spo0A (Spo0A∼P) levels influences cell development (9). The transition state is characterized by the initial activation of Spo0A at the end of exponential growth (6). An increase in Spo0A∼P levels and activation of the sigma factor σ^H^ directs cells into the sporulation program that is later driven by the compartment-specific master regulators of early sporulation σ^F^ and σ^E^, forespore and mother cell specific, respectively (10). The production of σ^F^ occurs before septation, which becomes active after septation in the forespore (6, 11, 12). The σ^F^ activation cascade includes SpoIIE, SpoIIAB, and SpoIIAA proteins, which are synthesized in the pre-divisional cell (13, 14). σ^F^ is inactive when bound with the anti-sigma factor SpoIIAB. SpoIIAA is an anti-anti-sigma factor that relieves the repression of SpoIIAB on σ^F^ activity(15). SpoIIE is a serine phosphatase(16) that catalyzes the dephosphorylation of SpoIIAA, thereby contributing to the forespore-specific activation of σ^F^ (11, 17). The expression of σ^F^-dependent genes is transcriptionally regulated by *rsfA*, a DNA-binding protein that blocks the premature appearance of σ^E^ (18). While σ^F^ is activated in the forespore, σ^E^ in the mother cell is activated after septation is complete (6). Pro-σ^E^ is synthesized before the asymmetric division from the Spo0A∼P dependent operon *spoIIG* (19). SpoIIGA is a protease that cleaves pro-σ^E^ and activates it (20, 21).

Upon activation of both sigma factors, entry into sporulation becomes irreversible (22, 23). Activation of the compartment-specific sigma factors σ^F^ and σ^E^ is initiated by asymmetric division of the cell, which represents stage I of sporulation (6). Morphologically, during this stage the chromosome is replicated and extended from one pole to the other, forming a structure called the axial filament (24). The formation of asymmetric septa occurs during stage II of sporulation. During stage III, the polar septum between the mother cell and the forespore curves around the forespore, leading to the engulfment of the forespore (4). The early engulfment stage is primarily driven by proteins synthesized in the mother cell, SpoIID, SpoIIM, SpoIIP, which are under the control of σ^E^ (25, 26). Activation of σ^G^ in the forespore, followed by activation of σ^K^ in the mother cell, signals the transition of the sporulation to stage IV (6, 27). Morphologically, the mother cell produces the proteinaceous coat that further assembles around the forespore (28, 29). Similarly to σ^F^, σ^G^ is held inactive by SpoIIAB in the pre-engulfment forespore, and the activation of σ^G^ requires its release (30, 31). Activation of σ^G^ in the forespore is followed by activation of the mother cell specific σ^K^ (6, 31). Transcription of *sigK* occurs in the mother cell from a σ^E^-dependent promoter (32). Similarly to σ^E^, σ^K^ appears initially as pro-σ^K^ approximately one hour before the process, depending on the gene expression in the prespore (32). The σ^G^ regulon includes *ssp* genes, which encode the small acid soluble proteins (SASPs), late sporulation genes, and genes that control germination (33, 34, 35, 36). The σ^K^ regulon directs transcription of 14 spore coat genes, including *cotD* and *spoVK* (37). Stage V of sporulation is characterized by the forespore depositing a protective exterior consisting of two layers of peptidoglycan, the primordial germ cell wall, and the cortex (38). This is followed by stage VI, which is the final maturation of the spore (3). The late stages of sporulation include spore DNA coating with SASPs, which protects spores from heat and UV radiation, and packaging of dipicolinic acid inside the spore core, to protect against heat and desiccation (39, 40). The mature spore is released during the final stage VII through the lysis of the mother cell (3, 6, 9).

The mature spore possesses phenomenal resistance properties, allowing for survival under extreme environmental conditions, such as exposure to heat, radiation, antibiotics, disinfectants, and lack of nutrients (41, 42). The structural protection of the spore chromosome from such insults is one reason for these properties, which is dependent upon the activity of the SASPs (33, 43, 44, 45). Previously, the histone-like protein HBsu was implicated to play a role in condensation of the spore chromosome, in addition to the SASPs (40). HBsu colocalizes with the SASPs on the DNA, in both the mother cell and spore, and counteracts SASP-mediated changes in persistence length and supercoiling (40). It was proposed that HBsu counteracting the SASP-mediated increase in persistence length might be required for chromosome packaging during sporulation. So far, it is known that HBsu binds non-specifically to DNA and regulates DNA compaction, replication initiation, recombination, and repair (46, 47, 48, 49, 50)

Our lab is interested in understanding the regulatory role of HBsu acetylation. N^Ɛ^-lysine acetylation is a ubiquitous regulatory post-translational modification (PTM), that can influence protein conformation and interactions with substrates, leading to change of function (51, 52, 53). Previously, we identified seven acetylation sites on HBsu and found that acetylation at some residues reduced the DNA binding affinity and regulated nucleoid compaction (54, 55). Furthermore, we investigated whether the acetylated state of HBsu can influence the process of sporulation and the resistance properties of mature spores (53). Using glutamine substitutions to mimic the acetylated state and arginine substitutions to mimic the unacetylated state, we found that certain mutants led to a reduction in sporulation frequency and resistance to heat, formaldehyde, and ultraviolet (UV) light (53). We proposed that acetylation at key residues of HBsu is required for proper sporulation; however, the exact underlying mechanism by which HBsu acetylation regulates this process is unknown.

We hypothesized that one mechanism by which acetylation of HBsu could influence sporulation is by the regulation of sporulation-specific gene expression. To test this hypothesis, we selected important sporulation-specific genes and regulatory factors that are turned on throughout the sporulation processes and examined expression by quantitative real-time PCR (qRT-PCR). We found that the *hbsK41Q* mutant was different from the other mutants, in that the expression of 17/25 genes analyzed were significantly upregulated at the later sporulation timepoints, whereas most of the other mutant strains exhibited the opposite phenotype. *hbsK3Q, hbsK37Q, hbsK75Q, hbsK80Q, hbsK86Q* strains had downregulated several important genes at T_4_. Based on these findings, we propose that acetylation at K41 plays a significant regulatory role during sporulation, which might be responsible for switching on or off gene expression as the cell moves from the early to late developmental stages.

## Results

### Selection of reporter genes and RT-qPCR assay development

We hypothesized that one way acetylation of HBsu influences the process of sporulation is through the regulation of gene expression. To test this idea, a quantitative real-time PCR (RT-qPCR) assay was developed and optimized. We examined four different house-keeping genes to serve as an internal control, *rpoA*, *rrnA-16S*, *rrnA-5S*, and *rpoD.* The efficiency of each set of primers (Table S2) was determined and *rpoD* was ultimately selected as the internal control for normalization of differing amounts of input cDNA. Based upon the expression patterns of sporulation-specific genes, 25 genes were selected for analysis (6, 56). The selected genes were representative of various early and late stages of sporulation. Samples were collected at 4 time points, T_0_-T_4_. The T_0_ time point corresponds to the beginning of sporulation (stages 0-I), when the cells “decide” to sporulate. The T_1_ time point corresponds to stages II-III of sporulation, characterized by significant changes in the cell morphology, including asymmetric septation and engulfment. The T_2_ time-point is later in the process and corresponds to stage IV of sporulation, while T_4_ corresponds to stage V. As the sporulation-specific sigma factors are important master regulators driving this process, various regulon members were included as reporters of sigma factor activity (Table S3, Fig. S1). Additionally, we analyzed the asynchronous cultures at the selected time-points to determine and confirm the staging of sporulation. As expected, there were cells at different stages concomitantly within the same population (Fig. S2).

### Acetylation at lysine 41 increases expression of the early sporulation genes

To examine the influence of acetylation, we utilized our collection of point mutations at the native *hbs* locus that mimic the acetylated state (glutamine substitutions (57)). The *hbsK41Q* mutant and wild-type strains were grown in DSM and analyzed throughout sporulation from T_0_-T_4_. First, we examined the activity of the housekeeping sigma factor σ^A^ and the stationary phase sigma factor σ^H^, which are responsible for the entry into the sporulation process, by examining the expression of the phosphorelay proteins (*kinA, kinB, spo0A, spo0B,* and *spo0F*). At T_0_ and T_1_ time points, representing early sporulation when these genes are likely important, we did not observe significant differences in expression compared to the wild type, except for *spo0F*, which was slightly elevated at1.35-fold, (p= 0.0243, Fig. 1). At T_2_, there was a 2.6-fold increase in *spo0A* expression. In contrast, at the T_4_ time-point there were significantly reduced levels of *spo0F*, *spo0A, kinA*, and *kinB* transcripts, but this is likely not biologically relevant as phosphorylated Spo0A is not required at this late stage.

**Figure 1:**
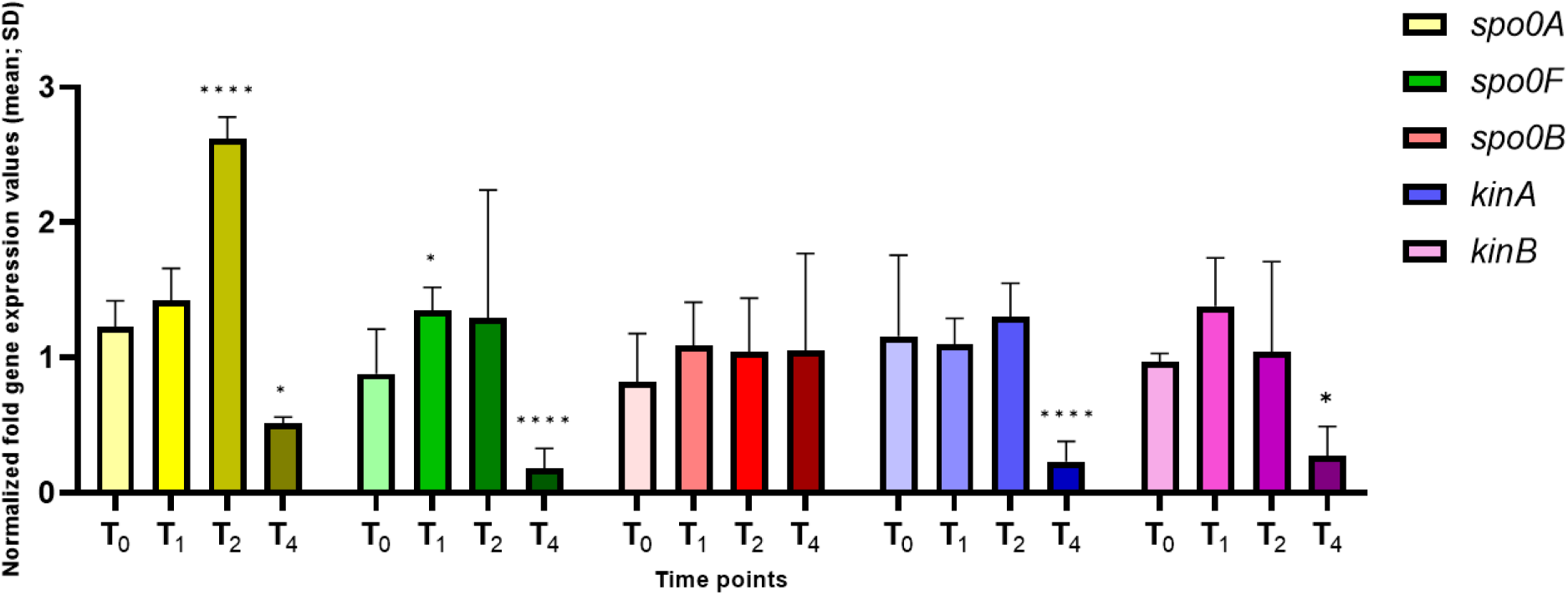
*Expression of the phosphorelay genes in the hbsK41Q mutant.* Wildtype and *hbsK41Q* mutant cells were grown in DSM for 6 hours and analyzed at time points T_0_-T_4_. Values displayed are the mean expression of technical duplicates from at least three biological replicates with the standard deviation (SD). The relative expression of the phosphorelay genes was similar to wildtype at T_0_, T_1_, and T_2_, except for *spo0F* (1.35-fold increase, p = 0.0243) at T_1_ and *spo0A* at T_2_ (2.62-fold increase, p< 0.000001). At T_4_, there was a statistically significant decrease in expression of *kinA, kinB, spo0A,* and *spo0F*. *p <0.05, ****p< 0.0001

Next, we examined the expression of genes that are dependent upon Spo0A∼P, including *spoIIE,* and the *spoIIA* and *spoIIG* operons, with the proteins required during stage II of sporulation (Fig. 2). Expression of these genes typically occurs before asymmetric septation and they are all involved in the activation of σ^F^. *spoIIAB* was overexpressed modestly at T_1_, with a 1.46-fold increase (p= 0.019), but an 18.4-fold (p= 0.0173) increase at T_2_. Despite these increases in transcription, a corresponding increase in protein levels was not observed (Fig. S3). Similarly, there was a 1.86-fold overexpression of *spoIIGA* at T_1_ (p= 0.0398) and 41.8-fold overexpression at T_2_ (p= 0.0502). *spoIIE* was significantly overexpressed 34.1-fold (p= 0.0094) at T_2_. At T_4_, *spoIIAB* and *spoIIE* were modestly decreased compared to wildtype, both 0.45-fold (p= 0.003 and 0.012, respectively). This suggests that the increase in *spo0A* transcription observed at T_2_ leads to more Spo0A∼P in the cell, which increases expression of Spo0A-dependent promoters, especially at T_2_.

**Figure 2:**
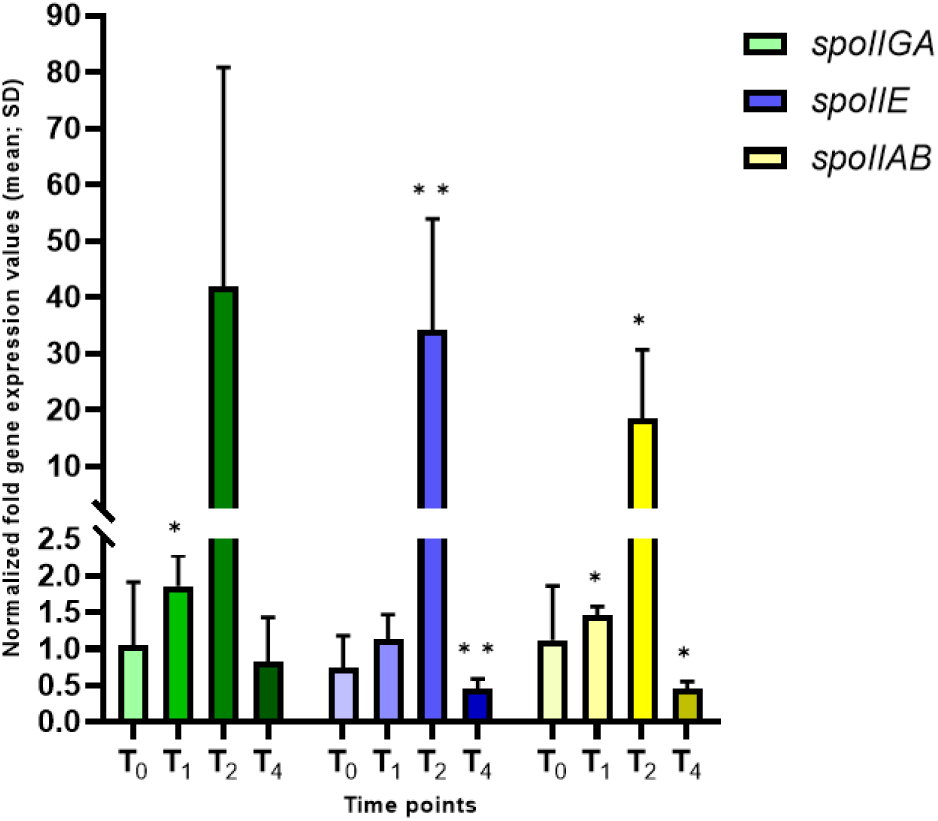
*Dynamics of Spo0A/σ^A^ and Spo0A/σ^H^ transcription over T_1_-T_4_ timepoints in the hbsK41Q mutant*. Wild-type and the *hbsK41Q* mutant cells were grown in DSM for 6 hours and analyzed at time points T_0_-T_4_. Values displayed are the mean expression of technical duplicates from at least three biological replicates with the standard deviation (SD). An increase in Spo0A∼P levels led to a modest upregulation of *spoIIGA* and *spoIIAB* at T_1_, and a large increase in expression at T_2_ (41.8-fold (p= 0.0502) and 18.4-fold (p= 0.017), respectively). *spoIIE* was significantly overexpressed 34.1-fold (p=0.0094) at T_2_. *p< 0.05, **p< 0.01

σ^F^ and σ^E^ are compartment-specific sigma factors that are active in the forespore and mother cell, respectively. Engulfment is the morphological process following σ^E^ activation, which occurs under the transcriptional control of both σ^F^ and σ^E^. The coordinated transcription between the distinct cellular compartments leads to the completion of engulfment, generating a double-membrane organelle within the mother cell. We found that *sigF* transcription was significantly increased at T_2_ (16.83-fold, p= 0.0073), and decreased at T_4_ (0.29-fold difference, p= 0.0001, Fig. 3). We analyzed expression levels of σ^F^ by Western blot, and found that in agreement with RT-qPCR, σ^F^ was overexpressed 6 and 7.6-fold at T_2_ and T_4_, respectively, even though transcription was reduced at T_4_ (Fig. 4). The increased protein levels, coupled with wild-type levels of the anti-sigma factor SpoIIAB (Fig. S3), implied that σ^F^ activity will also be increased, so σ^F^-dependent genes were analyzed, including *spoIIQ* and *sigG*. For *spoIIQ*, there was a 31.7-fold overexpression at T_2_, although not statistically significant (p= 0.0609), due to the presence of outliers among replicates. At T_2_ there was an overexpression of *sigG* (35.1-fold, p= 0.0157) and at T_4_ there was a 3.69-fold increase (p= 0.0029) in expression. In agreement, there was a 10.3-fold increase in protein at levels at T_4_ (Fig. 5). Together, these results suggest that overproduction of σ^F^ at T_2_ leads to inappropriately increased expression of the σ^F^ regulon.

**Figure 3:**
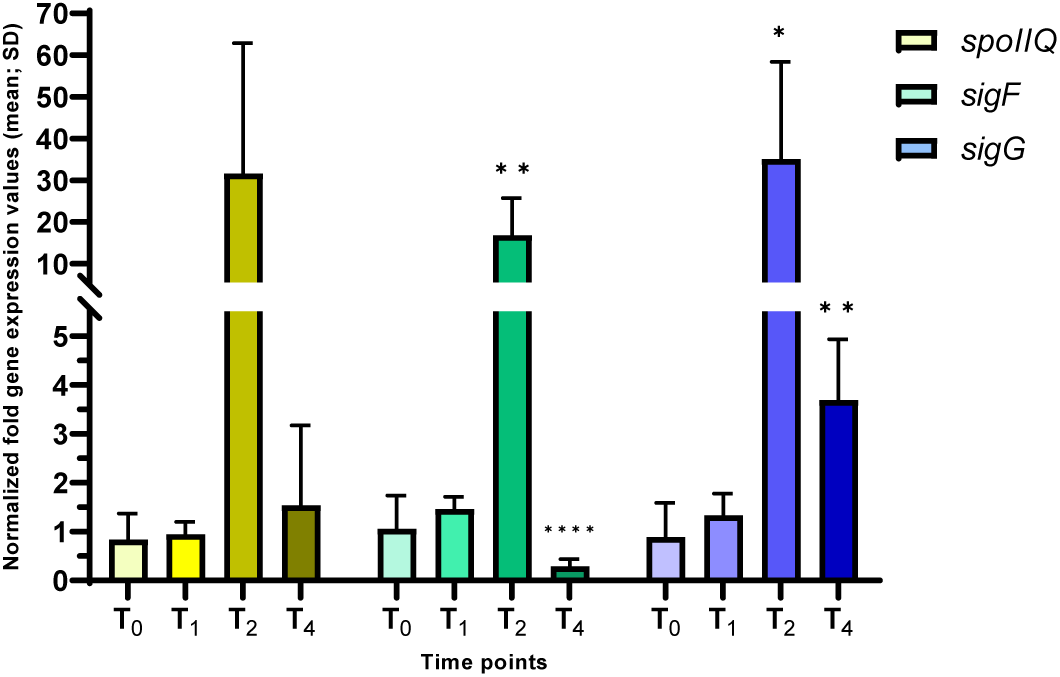
*σ^F^ activity in the hbsK41Q strain.* Wildtype and the *hbsK41Q* mutant were grown in DSM for 6 hours and analyzed at time points T_0_-T_4_. Values displayed are the mean expression of technical duplicates from at least three biological replicates with the standard deviation (SD). The forespore-specific genes *sigF* and *sigG* were significantly upregulated 16.82-fold and 35.12-fold, respectively, at T_2_ (p = 0.0073 and p = 0.0157, respectively). At T_4_, *sigG* expression was increased 3.69-fold (p = 0.0029) and *sigF* expression was 0.29-fold (p = 0.0001) compared to wildtype. *spoIIQ* expression mirrored the *sigF* expression pattern but was not statistically significant due to high variability among replicates. *p< 0.05. **p< 0.01, ****p <0.0001

**Figure 4.**
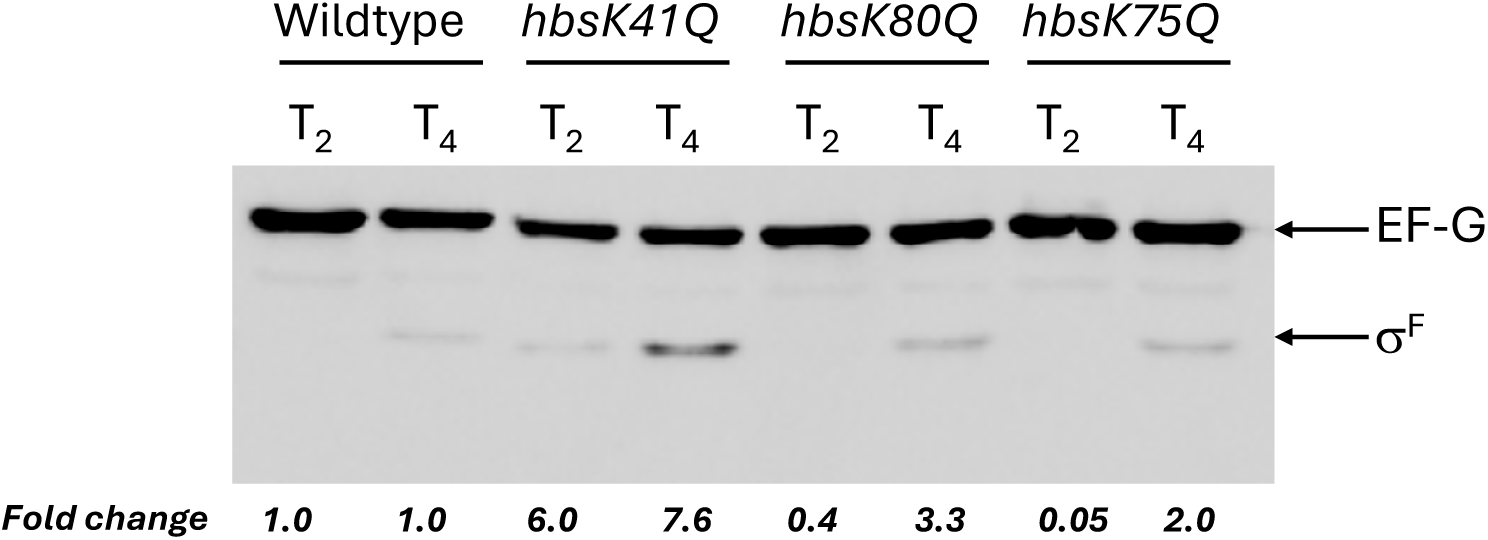
*σ^F^ levels in acetylation mutants.* Wild-type, *hbsK41Q*, *hbsK80Q, hbsK75Q* cells were grown to T_2_ and T_4_ in sporulation media and lysates prepared as described in Materials and Methods. Equal amounts of protein were loaded and were probed with anti-σ^F^ and anti-EF-G antibodies. EF-G was included as a loading control. Non-labeled bands represent cross-reacting bands. All western blots were repeated three independent times, and a representative blot is shown. Band Intensities were quantified using ImageJ, and fold changes were calculated by normalizing to EF-G and mutants compared to wildtype.

**Figure 5:**
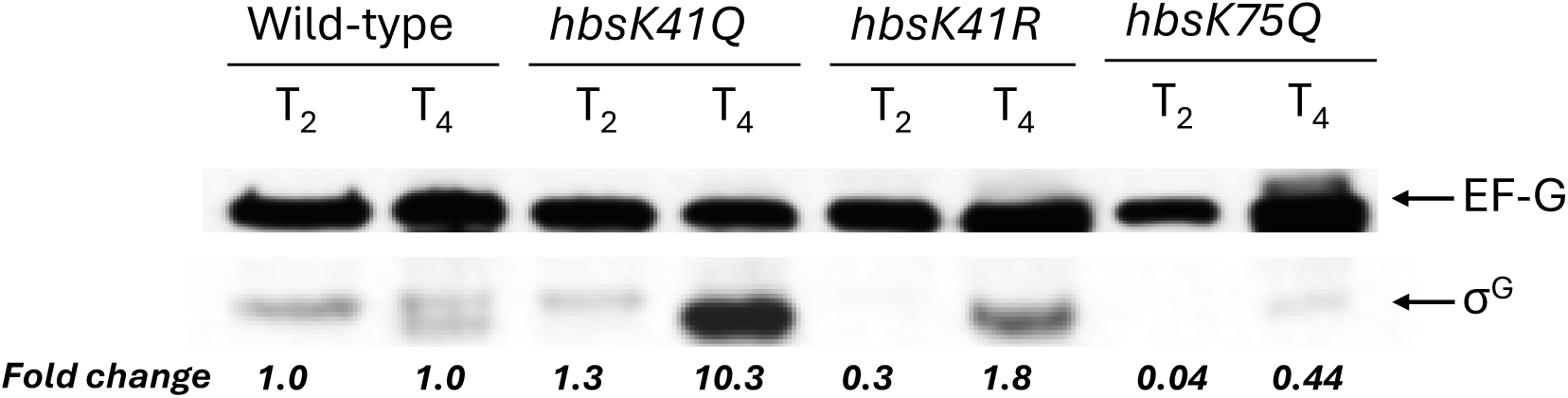
*σ^G^ levels in acetylation and deacetylation mutants.* Wild-type, *hbsK41Q*, *hbsK41R, hbsK75Q* cells were grown to T_2_ and T_4_ in sporulation media and lysates prepared as described in Materials and Methods. Equal amounts of protein were loaded and were probed with anti-σ^G^ and anti-EF-G antibodies. EF-G was included as a loading control. (A) Non-labeled bands represent cross-reacting bands. All western blots were repeated three independent times, and a representative blot is shown. The EF-G and σ^G^ blots are from the same gel with different exposure times during development. The σ^G^ signal is weaker and requires a longer exposure than EF-G, which becomes saturated. The complete, unmodified blots taken at the different exposure times are provided in Figure S4. Band Intensities were quantified using ImageJ, and fold changes were calculated by normalizing to EF-G and mutants compared to wildtype.

To assay the mother-cell specific σ^E^ activity, *asnO*, *spoIID, spoIVCA,* and *spoVK* were analyzed. There was a large increase in *spoIID* levels (26.58-fold, p= 0.0629), which is only dependent upon σ^E^ for activation, suggesting increased sigma factor activity. *asnO* was significantly increased across the entire course, with a 1.79, 2.83, and 65.98-fold increase at T_1_, T_2_, and T_4_, respectively (p= 0.0218, 0.0166, and 0.0432, respectively). *spoVK* and *spoIVCA* were both overexpressed at the later time points, while not statistically significant (Fig. 6). These results suggest that σ^E^ activity is also increased when K41 is acetylated.

**Figure 6:**
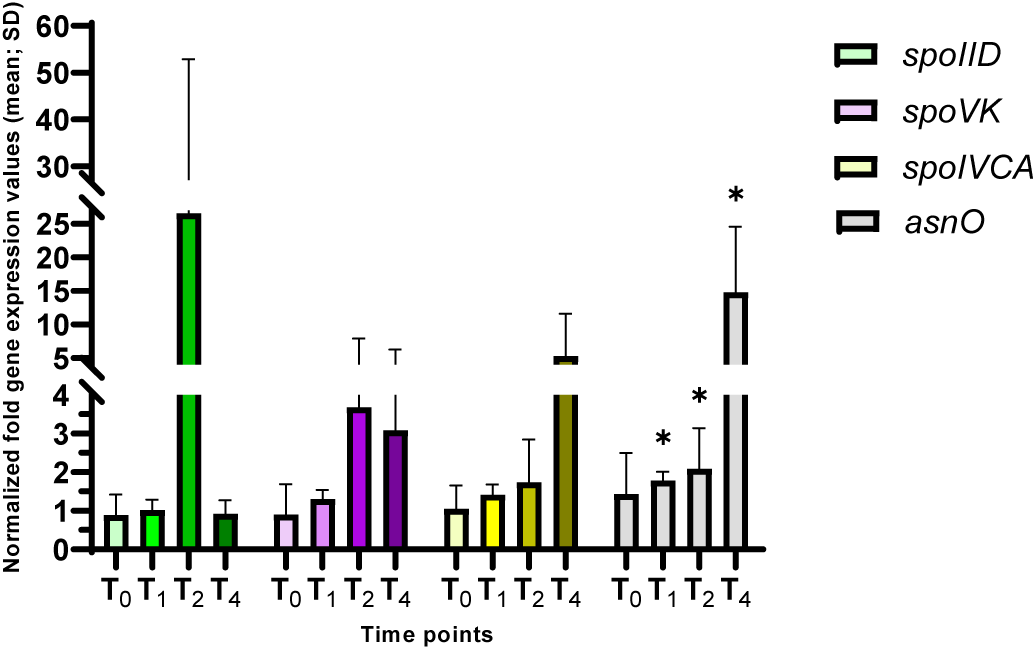
*σ^E^ activity in hbsK41Q strains.* Wildtype and the *hbsK41Q* mutant were grown in DSM for 6 hours and analyzed at time points T_0_-T_4_. Values displayed are the mean expression of technical duplicates from at least three biological replicates with the standard deviation (SD). *spoIVCA,* a protein that processes pre-σ^K^, and *spoVK*, which is involved in cortex formation, were increased at the later time points, although not significantly. The asparagine synthetase gene, *asnO* was overexpressed from T_1_ to T_4_ at 1.79-fold (p= 0.0218), 2.83-fold (p= 0.0166), and 72.6–fold (p= 0.0432), respectively. There was a large increase in expression of *spoIID* at T_2_, which did not reach significance due to the presence of outliers (26.58-fold, p= 0.0629). *p< 0.05

### Acetylation at lysine 41 increases expression of the late sporulation genes

The completion of engulfment stage dramatically changes the transcriptional program within both compartments, with the activation of σ^G^ in the forespore. To assay σ^G^ activity, we examined the expression of *spoVAA*, *gpr*, *spoVT* and *sspB* (Fig. 7). At the T_4_ time point *spoVT* was overexpressed 7.87-fold (p= 0.0193) and *sspB*, which is solely dependent on σ^G^ for expression, was overexpressed at 2.72-fold, 4.47-fold, and 61.6-fold at T_1_, T_2_, and T_4_ (p= 0.0792, 0.3979 and 0.2233, respectively). The presence of a large outlier in one replicate (>120-fold increase, compared to ∼30-fold for all replicates) is the reason statistical significance was not reached T_4_, despite the large increase in expression. *spoVAA* was overexpressed at T_2_ (6.59-fold, p= 0.0013) and T_4_ (26.79-fold, p= 0.0126), whereas *gpr* did not exhibit large differences in expression.

**Figure 7:**
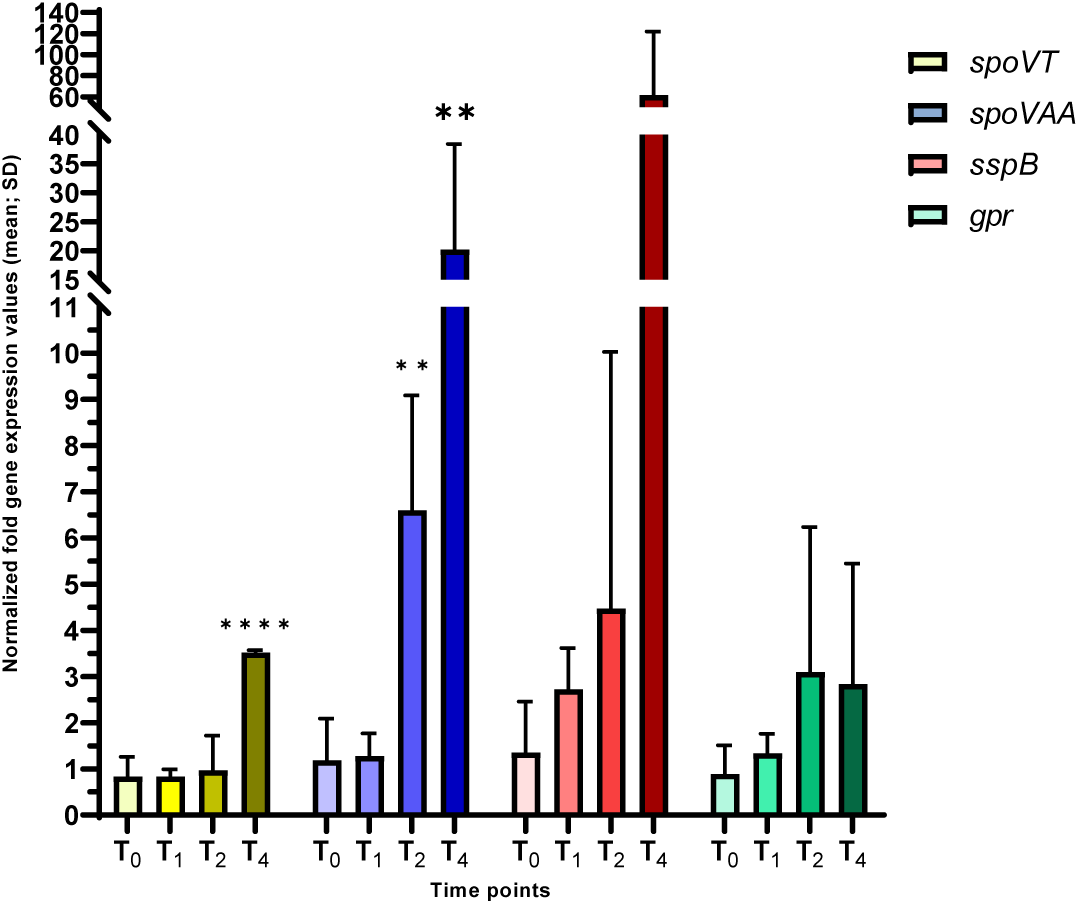
*σ^G^ activity in the hbsK41Q mutant.* Wildtype and the *hbsK41Q* mutant were grown in DSM for 6 hours and analyzed at time points T_0_-T_4_. Values displayed are the mean expression of technical duplicates from at least three biological replicates with the standard deviation (SD). The σ^G^ regulon members were significantly overexpressed at T_4_. The transcriptional regulator *spoVT*, was 7.87-fold increased at T_4_ (p= 0.0193). *spoVAA*, which encodes dipicolinic acid uptake protein, was overexpressed at T_2_ (6.59-fold, p= 0.0013) and T_4_ (26.8-fold, p= 0.0126). *sspB*, which encodes a ß-type small acid-soluble protein, was overexpressed at T_4_, which was not statistically significance due to high variability among replicates. *gpr* did not show any statistically significant differences at any time point. * p< 0.01, ****p< 0.0001

To analyze the late stages of sporulation, we examined the expression of the coat proteins. The expression of *cotE* and *cotH* is under the control of σ^E^ and σ^K^, and *cotD* is under the control of only σ^K^. We observed a significant increase in the expression level of *cotE* (19.9-fold, p= 0.0367) and *cotH* (18.3-fold, p= 0.0354) at T_4_ (Fig. 8). *cotD* and *lipC* expression were not different from wildtype, suggesting that σ^K^ activity was not altered. The large increase in *cotE* and *cotH* expression likely resulted from the increase in σ^E^ activity. As we observed a gradual increase in the expression of *spoIVCA*, which encodes a protein essential for posttranslational processing of pre-σ^K^, we examined σ^K^ levels by western blot (Fig. 9). σ^K^ was overproduced at T_4_ compared to wildtype. σ^K^ is produced as a longer, inactive form that must be cleaved N-terminally for activation and both bands were detected. The higher band is the inactive protein before cleavage, and the smaller band represents the active protein. σ^K^ is overproduced at T_4_ compared to wildtype, with significantly more processing (92% compared to 56%).

**Figure 8.**
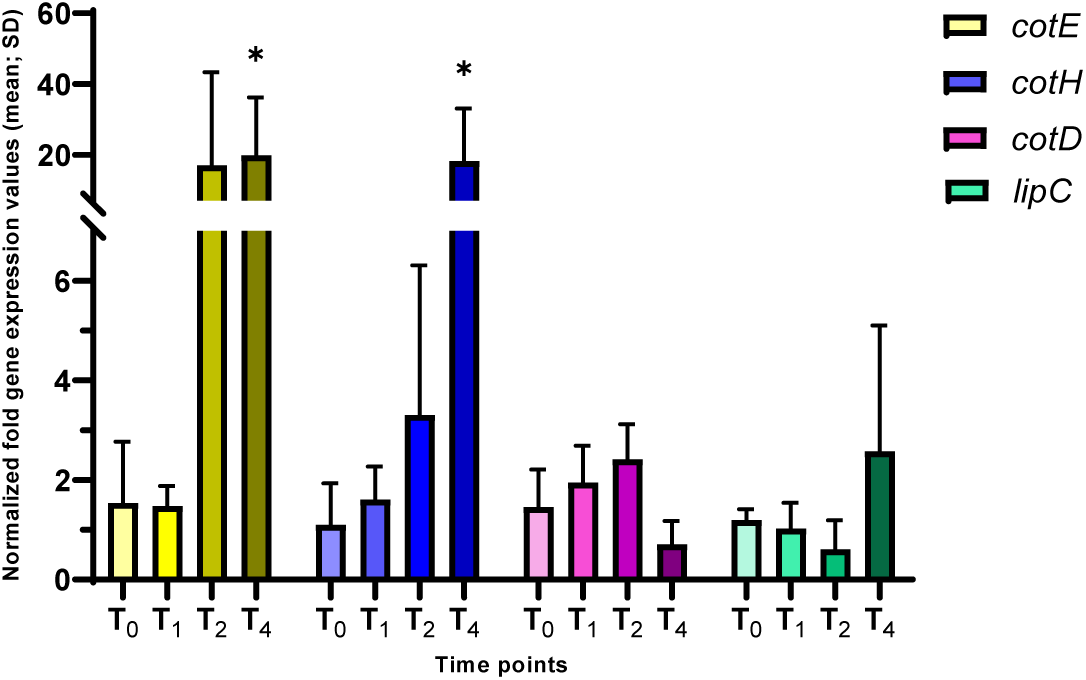
*σ^K^ activity in hbsK41Q.* Wildtype and the *hbsK41Q* mutant were grown in DSM for 6 hours and analyzed at time points T_0_-T_4_. Values displayed are the mean expression of technical duplicates from at least three biological replicates with the standard deviation (SD). The mother-cell-specific genes involved in coat formation were significantly overexpressed at T_4_, with *cotE* at 19.9-fold (p= 0.0367) and *cotH* at 18.3 (p= 0.0354). There were no significant differences in expression for *cotD* and *lipC*, which are exclusively dependent upon σ^K^ for transcription. * p< 0.05

**Figure 9.**
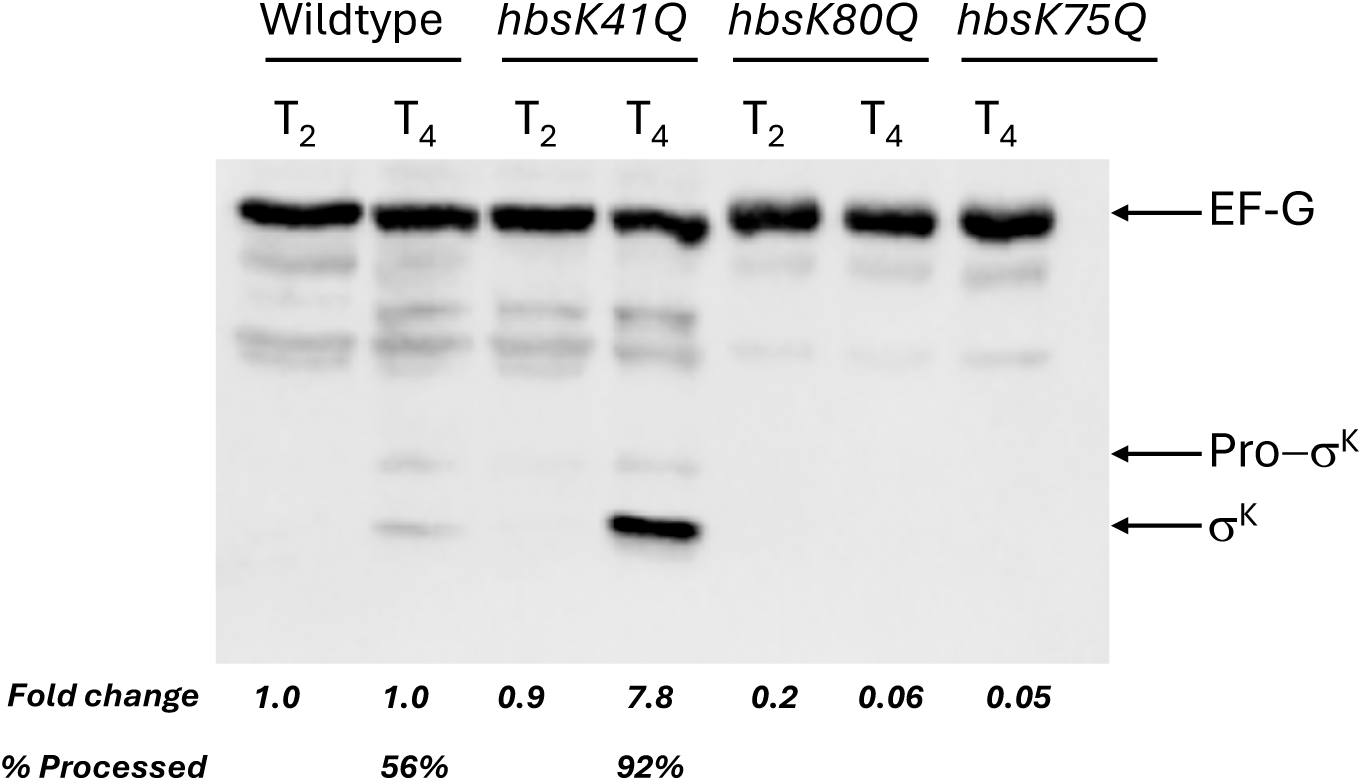
*σ^K^ levels in acetylation mutants.* Wild-type, *hbsK41Q*, *hbsK80Q, hbsK75Q* cells were grown to T_2_ and T_4_ in sporulation media and lysates prepared as described in Materials and Methods. Equal amounts of protein were loaded and were probed with anti-σ^K^ and anti-EFG antibodies. EFG was included as a loading control. Non-labeled bands represent cross-reacting bands. (A) σ^K^ is detected in two forms: inactive precursor pro-σ^K^ (top band) and activated σ^K^ (bottom band). All western blots were repeated three independent times, and a representative blot is shown. Band Intensities were quantified using ImageJ, and fold changes were calculated by normalizing to EF-G and mutants compared to wildtype. The values for full length σ^K^ are shown.

### Deacetylation of lysine 41 does not induce gene expression

Since we observed many changes associated with the *hbsK41Q* mutants, we analyzed the opposite, deacetylated form with the *hbsK41R* mutant strain. At T_2_, for most genes analyzed, similar levels to wildtype were observed (Table 2). *asnO, spo0F, spoVK* and *spoVG* were increased 2-to 6-fold in the *hbsK41Q* strain but displayed reduced expression in the *hbsK41R* strain. This suggests that many of these genes may be directly regulated by HBsu and acetylation at K41 is required for gene expression at T_2_. At T_4_, roughly one third of the transcripts that were significantly increased in the *hbsK41Q* strain were altered to the same extent in the *hbsK41R* strain. For example, *cotE* and *cotH* were both highly overexpressed in the *hbsK41Q* mutant (19.9-fold, p= 0.0397, and 18.3-fold, p= 0.0354, respectively) and were overexpressed to a similar extent in the *hbsK41R* mutant (27.4-fold, p= 0.0001, and 26.8-fold, p= 0.0008, respectively). This might be due to the influence of other transcription factors and suggests that HBsu K41 acetylation is not the primary regulatory determinant for expression of these genes at T_4_.

**Table 1.** Strains used in this study.

| *B. subtilis* strain number | Genotype* | References |
| --- | --- | --- |
| BD630 | *his leu8 metB5* | Laboratory strain |
| BD7493 | *hbsK86Q* | (57) |
| BD8119 | *hbsK37Q* | (57) |
| BD8147 | *hbsK41Q* | (57) |
| BD8148 | *hbsK41R* | (57) |
| BD8219 | *hbsK18Q* | (57) |
| BD8398 | *hbsK75Q* | (57) |
| BD8576 | *hbsK80Q* | (57) |
| BD8577 | *hbsK3Q* | (57) |

\*All strains are in the BD630 background

**Table 2.**
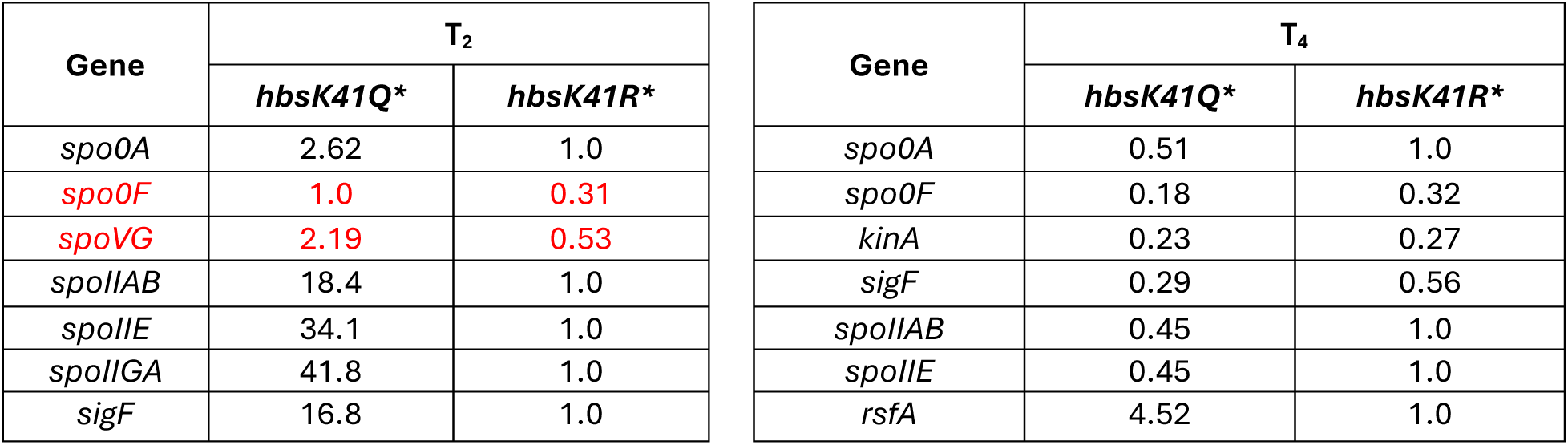
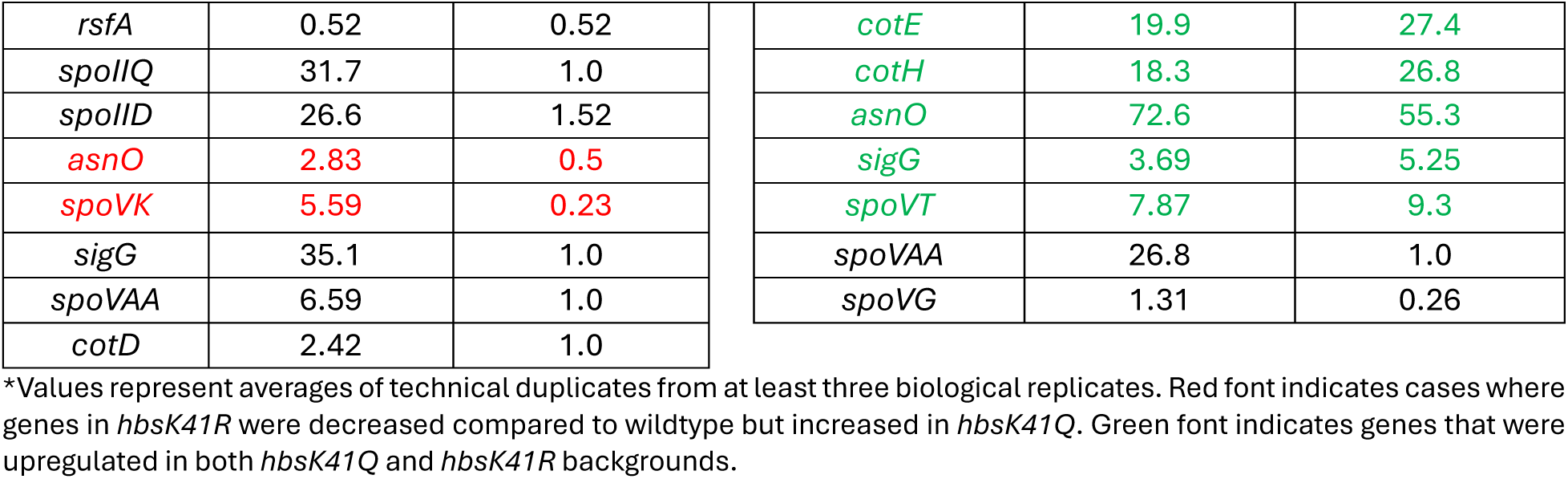
Comparison of gene expression between *hbsK41Q* and *hbsK41R* mutants at T_2_ and T_4_.

| Gene | $T_2$ | |
| --- | --- | --- |
|  | *hbsK41Q* * | *hbsK41R* * |
| *spo0A* | 2.62 | 1.0 |
| *spo0F* | 1.0 | 0.31 |
| *spoVG* | 2.19 | 0.53 |
| *spolIAB* | 18.4 | 1.0 |
| *spolIE* | 34.1 | 1.0 |
| *spolIGA* | 41.8 | 1.0 |
| *sigF* | 16.8 | 1.0 |

| Gene | $T_4$ | |
| --- | --- | --- |
|  | *hbsK41Q* * | *hbsK41R* * |
| *spo0A* | 0.51 | 1.0 |
| *spo0F* | 0.18 | 0.32 |
| *kinA* | 0.23 | 0.27 |
| *sigF* | 0.29 | 0.56 |
| *spolIAB* | 0.45 | 1.0 |
| *spolIE* | 0.45 | 1.0 |
| *rsfA* | 4.52 | 1.0 |

|  |  |  |  |  |  |
| --- | --- | --- | --- | --- | --- |
| *rsfA* | 0.52 | 0.52 | *cotE* | 19.9 | 27.4 |
| *spolIQ* | 31.7 | 1.0 | *cotH* | 18.3 | 26.8 |
| *spolID* | 26.6 | 1.52 | *asnO* | 72.6 | 55.3 |
| *asnO* | 2.83 | 0.5 | *sigG* | 3.69 | 5.25 |
| *spoVK* | 5.59 | 0.23 | *spoVT* | 7.87 | 9.3 |
| *sigG* | 35.1 | 1.0 | *spoVAA* | 26.8 | 1.0 |
| *spoVAA* | 6.59 | 1.0 | *spoVG* | 1.31 | 0.26 |
| *cotD* | 2.42 | 1.0 |  |  |  |

\*Values represent averages of technical duplicates from at least three biological replicates. Red font indicates cases where genes in *hbsK41R* were decreased compared to wildtype but increased in *hbsK41Q*. Green font indicates genes that were upregulated in both *hbsK41Q* and *hbsK41R* backgrounds.

### The acetylation state of HBsu at lysine 41 increases sporulation frequency at inappropriate times

Since most important sporulation genes were overexpressed and sigma factor activity was increased, we hypothesized that *hbsK41Q* strains sporulate earlier than the wildtype. To test this idea, we examined sporulation frequencies at T_0_ (Fig. S5), a time when the decision to enter sporulation usually occurs, and few or no intact spores should be observed. At T_0_, the wild-type and *hbsK41R* strains sporulated at a very low frequency of 0.04% and 0.07%, respectively, as expected (Fig. S4). However, there was a 25-fold increase in sporulation frequency of *hbsK41Q* strains (1%), which agrees with the idea that increased gene expression leads to earlier sporulation. Given that *hbsK41Q* is more susceptible to heat than the wildtype, and the sporulation frequency assay utilizes a 30-minute heat exposure to eliminate vegetative cells, we are likely underestimating the actual sporulation frequency.

### Acetylation of K3, K37, K75, K80, and K86 leads to a decrease in gene expression of key sporulation effectors

Previously, we found that significant reductions in sporulation frequency if the acetylation status was altered at K3, K18, K37, K75, K80, and K86(53). For each of these sites, we observed a reduction in the expression of key genes throughout the sporulation cycle in acetylation mutants, exactly opposite to that seen in the *hbsK41Q* strain. In the *hbsK3Q* strain at T_4_, the expression of two key late sporulation proteins, *cotD* and *sspB* were significantly reduced (0.32-fold, p= 0.0086, and 0.34-fold, p= 0.0223, respectively, Fig. 10). The *hbsK3Q* mutant has a reduced sporulation frequency, higher susceptibility to heat and formaldehyde exposure (53). Together, our data suggests when K3Q is acetylated, the late sporulation genes are not turned on at the proper time, which leads to the decreased heat and chemical stress resistance. Similarly, *cotD* and *sspB* were significantly reduced at the T_4_ in the *hbsK37Q* mutant (Fig. S6). The *hbsK37Q* strain was significantly more susceptible to heat exposure, with a survival rate below 10% after 30 minutes (53). *hbsK86Q* spores had a reduction in sporulation frequency (<50%) and were significantly more susceptible to heat exposure(53). There was a significant decrease in *spoIID* expression at T_4_ (Fig. S7A). There was also a significant decrease in the late sporulation genes *cotD* and *sspB* at T_4_ (Fig. S7B). This suggests that the acetylation of K3 K37, and K86 turn off gene expression of late sporulation genes. For the *hbsK3Q, hbsK37Q* and *hbsK86Q* mutants, the reduction in sporulation frequency observed might be completely attributed to the decreased wet-heat survival, as the gene expression program is largely intact.

**Figure 10:**
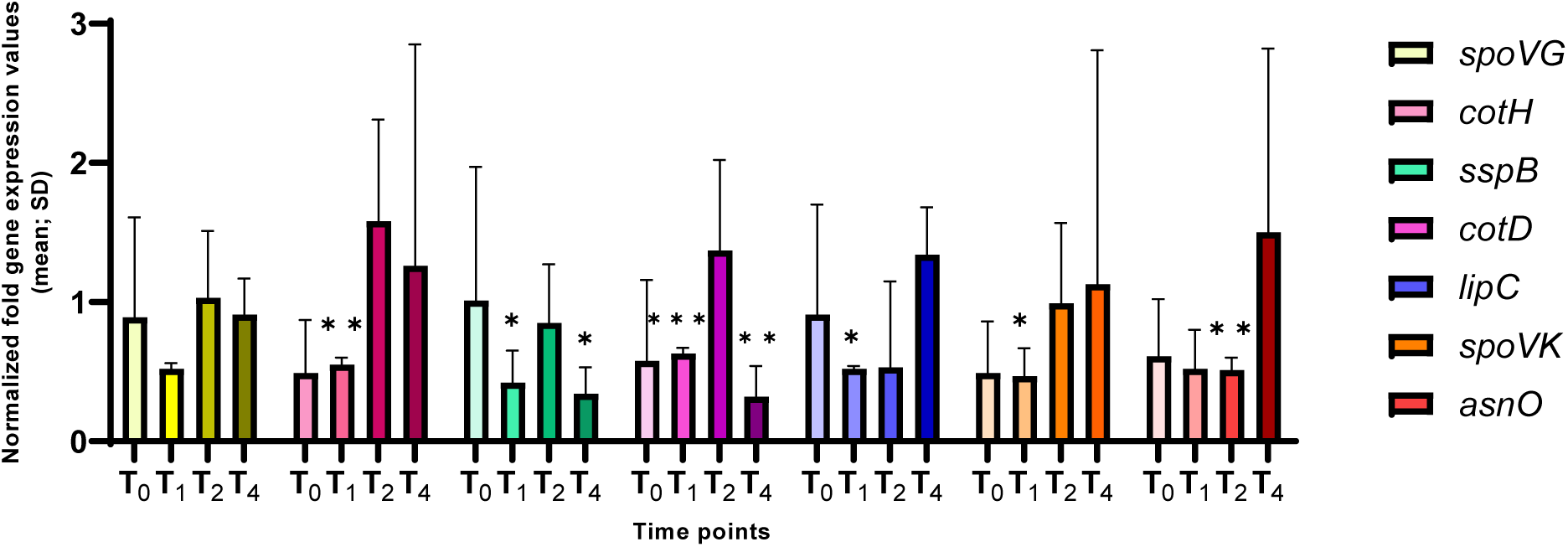
*Altered gene expression in hbsK3Q mutants*. Wildtype and the *hbsK3Q* mutant were grown in DSM for 6 hours and analyzed at time points T_0_-T_4_. Values displayed are the mean expression of technical duplicates from at least three biological replicates with the standard deviation (SD). The expression of the mother-cell specific *spoVK* (cortex assembly), *cotH* (coat assembly), and *lipC* (involved in germination) were reduced at T_1_. The late sporulation genes are not expressed at T_1_, so this is likely not biologically significant. *asnO* was ∼0.5-fold downregulated from T_0_-T_2_. The reduction in expression of *cotD* (coat protein) and *sspB* (ß-type small acid-soluble protein) to below 0.5-fold at T_4_ might explain the higher susceptibility to heat and formaldehyde in this mutant. *p< 0.05, **p< 0.01, ***p< 0.001

The *hbsK75Q* mutants have a reduced sporulation frequency, but the resulting spores were resistant to heat and UV stress. The only noted defect was reduced survival to formaldehyde after 40 minutes of exposure (53). Some early sporulation genes were modestly reduced, including *kinA, sigF, spo0A, spoIIQ*, and *spoIIAB* (Fig. 11A), with many of the significant effects observed at T_4_, when these genes would not be expected to be important. These changes likely do not explain the observed defects in sporulation frequency in this mutant. σ^F^ levels were reduced by about 50% at T_2_ compared to the wildtype (Fig. 4), while σ^G^ was reduced to 0.04 and 0.4 at T_2_ and T_4_, respectively (Fig. 5). σ^K^ levels were also reduced at T_4_, to 0.05-fold compared to wildtype (Fig. 9). In addition, some late sporulation genes were significantly reduced at T_4_, including *cotE*, *cotH*, and *sspB* (Fig. 11B and C), which might explain reduced survival to formaldehyde.

**Figure 11:**
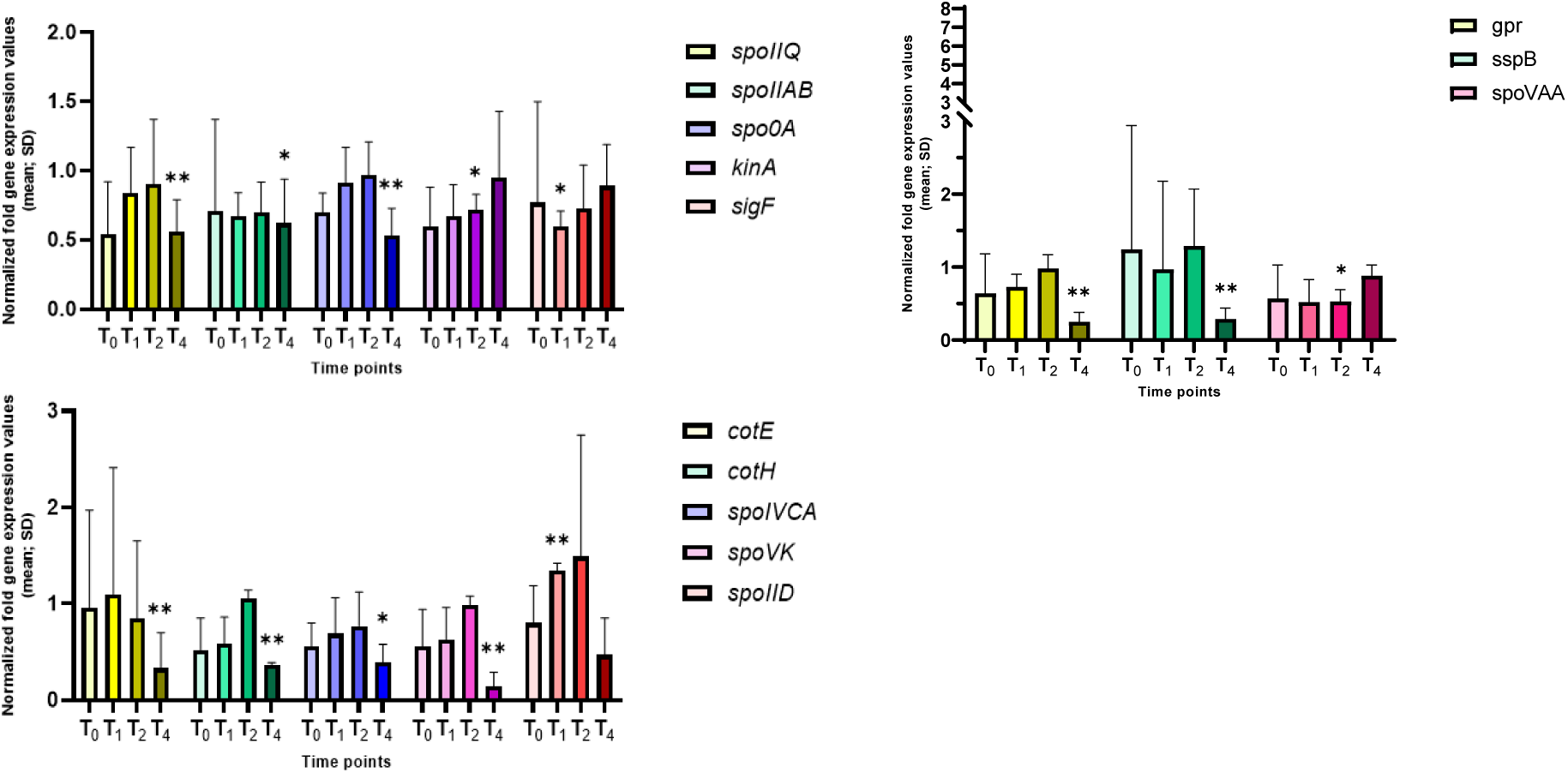
*Altered gene expression in hbsK75Q mutants.* Wildtype and the *hbsK75Q* mutant were grown in DSM for 6 hours and analyzed at time points T_0_-T_4_. Values displayed are the mean expression of technical duplicates from at least three biological replicates with the standard deviation (SD). (A) The early sporulation genes dependent upon Spo0A and σ^H^, or σ^F^ for activation, displayed modestly reduced gene expression at different times throughout sporulation. Many late sporulation genes dependent upon σ^K^ (B) or σ^G^ (C) were significantly reduced at T_4_. *p< 0.05, **p< 0.01

The *hbsK80Q* mutant had the second most alterations in gene expression, with most genes, especially the mother-cell specific ones, significantly downregulated. Previously, we found that the sporulation frequency of *hbsK80Q* strain was decreased by more than 50%, and that they were more susceptible to formaldehyde and UV radiation(53). Some forespore-specific early sporulation genes were downregulated, often at the T_1_ and T_2_ time points (i.e., *kinA, sigF, spoIIQ, spoIIAB, spoIIE*), which could, at least in part, explain the reduction in sporulation frequency of this mutant (Fig. 12A). There were also statistically significant decreases in σ^E^-and σ^K^-dependent genes. Most of these genes had decreased expression by more than 2-fold compared to the wildtype at the T_4_ time point (Figs. 12B and C). Given that these late genes are involved in forming protective layers of the spores, the observed decrease in expression might explain the higher susceptibility to chemical and UV stresses. As there were multiple under-expressed genes, we next examined the levels of the sigma factors. There were no differences in σ^G^ or σ^F^ levels compared to the wildtype (Figs. 4 and 5). However, we were unable to detect σ^K^ (Fig. 9), which might explain the reduction in the transcription of *sspB* and the *cot* genes. As acetylation of HBsu at K80 leads to decreased gene expression, this suggests that deacetylation of K80 is required for proper late gene expression.

**Figure 12:**
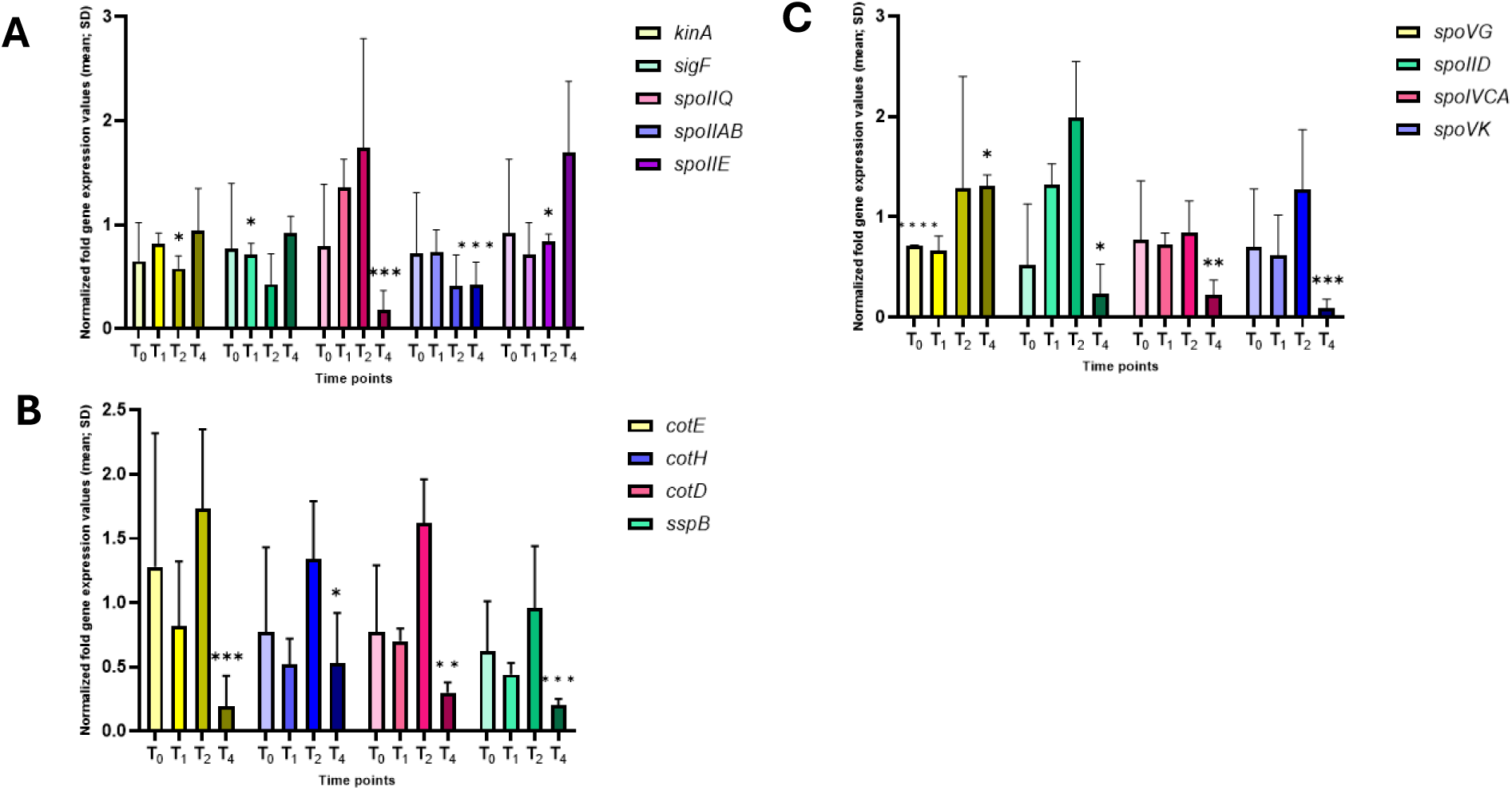
*Altered gene expression in hbsK80Q mutants.* Wildtype and the *hbsK80Q* mutant were grown in DSM for 6 hours and analyzed at time points T_0_-T_4_. Values displayed are the mean expression of technical duplicates from at least three biological replicates with the standard deviation (SD). (A) For the early sporulation genes, some significant downregulated changes were identified, but this occurred mostly after T_0_ and T_1_, which is these genes are required. (B) There was a significant reduction in expression of σ^K^-(B) and σ^G^-dependent (C) mother-cell specific reporters at T_4_. *p< 0.05, **p< 0.01, ***p< 0.001

## Discussion

HBsu is a histone-like protein essential for cell viability, normal growth, and development of *B. subtilis* (49, 58). Similarly to eukaryotic histones, the acetylation of HBsu decondenses chromosome by reducing its DNA-binding affinity (57, 58, 59). Previously, we found that HBsu contains seven acetylation sites (54). Given HBsu’s vital role in *B. subtilis* chromosome dynamics and DNA compaction, it is not surprising that HBsu was found in mature spores, at levels equal to that seen in vegetative cells (40). Recently, we found that certain acetylation sites on HBsu were involved in the regulation of the process of sporulation (53). The sporulation frequency of most of the acetyl-mimicking mutants were significantly reduced, except for the *hbsK41Q* mutant(53). In addition, we found that high levels of acetylation at K75 and K80 and some intermediate levels of acetylation at K3, K41 and K86 are required for the proper protection of spores from wet heat exposure(53). We also found that acetylation at K37, K41, K75, and K80 were important for survival of chemical exposure. This work did not examine the underlying mechanism as to how HBsu acetylation influences sporulation. Based upon the known functions of HBsu, there were two likely possibilities. First, as HBsu plays a role in chromosome organization and dynamics, it might be important for axial filament formation or compaction of the chromosome into the mature spore. HBsu colocalizes with the SASPs in the mature spore and modulates SASP-mediated increases in DNA-persistence length and supercoiling.(40) it is possible that HBsu acetylation modulates these activities. The second possibility is that HBsu influences gene expression. These possibilities are not mutually exclusive.

Here we addressed the latter possibility and examined gene expression throughout the process of sporulation to assess the impact of the acetylated forms of HBsu. Overall, all the mutants, except for *hbsK18Q*, had significant changes in gene expression. Acetylation at some sites changed gene expression dramatically, like *hbsK41Q* and *hbsK80Q*, although in opposite directions. These effects might have been amplified by altering the activity global regulators like the sigma factors. The most dramatic changes in the gene expression were observed in the *hbsK41Q* mutant. Many early sporulation genes (*kinA, spo0A, sigF, spoIIE, spoIIAB, spoIIQ, spoIIGA, spoIID*) and late sporulation genes (*spoVAA, sigG, cotD, spoVK, asnO*) were overexpressed at T_2_ (Figs. 1-3 and 6-8, respectively). Even though we examined an asynchronous population, we do not expect to find significant expression of the early sporulation genes at T_2_ because most cells have already transitioned into the sporulation program (Fig. S2). At T_4_, the late sporulation genes were highly upregulated (*cotE, cotH, sigG, spoVAA, spoVT*), possibly due to overexpression of sigma factors and transcription factors at earlier time points (Figs. 7 and 8). The opposite *hbsK41R* mutant did not lead to overexpression of any genes at T_2_ (Table 2), suggesting that at T_2_, acetylation at K41 activates gene expression. At T_4_, about a third of the genes were increased to a similar extent to that seen in the *hbsK41Q* strain (Table 2). It is important to note that the mutant strains represent a state with 100% acetylation or deacetylation at the specific site, whereas in nature, the process is continuous, and it is unknown if this state ever exists. Taken together, some acetylation level might be required during late sporulation at specific genes, but our data suggests that K41 should be mostly deacetylated for proper gene expression. We do not know if in wild-type cells, K41 is acetylated early in sporulation to drive gene expression and becomes deacetylated at T_2_ to turn off expression or if under normal conditions, acetylation occurs at T_2_ to activate gene expression. In either case, it appears that acetylation at K41 serves as an “on-off” switch and may drive the transition between early and late stages of sporulation. We are currently developing a mass spectrometry-based proteomics assay to measure the *in vivo* stoichiometry of acetylation of HBsu throughout the sporulation process, which will address these possibilities. One way of controlling this transition might be through the regulation of the expression and activity of σ^F^, σ^G^, and σ^K^. We found that all three of these sigma factors were overproduced at T_4_ (Figs. 4, 5, and 9), which likely disrupts the timing of the genetic program, and consequently leads to the loss of resistance properties of mature spores. In agreement, we found that *hbsK41Q* cells had increased levels of sporulation during vegetative growth, suggesting they enter the sporulation program early, even without appropriate signals (Fig. S5). The K41 site appears to be important for stationary phase development. From our original characterization of acetylation in HBsu, we found that acetylation of K41 was the only site with significantly increased abundance in stationary phase compared to exponential phase in glucose minimal media (54). We also found that *hbsK41Q* mutants had larger, expanded nucleoids in response to drug challenge, which is opposite to the normal wild-type response, which is to compact the nucleoid (60). In the same study, we observed 2.26-fold increase in the formation of persisters in the *hbsK41Q* background, and a faster recovery from the persistent state, compared to the wild-type (60). We proposed that the acetylation of K41 decondenses the chromosome to allow for cells to “escape” from the persistent state and resume growth. During sporulation, the acetylation of K41 likely triggers a transition between early and late sporulation, driven by altered gene expression. Given these findings, acetylation of K41 might also regulate gene expression to allow cells to escape from a drug-induced persistent state. This possibility has yet to be determined.

For all the other mutants, except for *hbsK18Q*, there was a downregulation of sporulation-specific genes. Acetylation of *hbsK18Q* likely does not influence gene expression, which cannot explain the observed reduction in sporulation frequency previously observed (53). It is still possible that acetylation at K18 influences chromosome dynamics or compaction of DNA into the mature spore, which will be addressed as part of a future study. The most significant changes were found in *hbsK80Q* and *hbsK75Q* (Figs. 11 and 12). The effects were largest at the later time points, especially T_4_. In the *hbsK80Q* strain, many early and late sporulation-specific genes were downregulated more than 2-fold compared to the wildtype (*cotE, spoIIQ, spoIIAB, spoIVCA, spoVK, cotD, spoIID, sspB*). A similar picture was observed in *hbsK75Q* with downregulation of *cotE, cotH, spoIIQ, spoIIAB, spoIVCA* at T_4_. This could be due to the proximity of K75 and K80 residues within the structure of the HBsu molecule. Acetylation of either site may turn off the expression of the same genes. In support of this idea, both mutants were resistant to heat but sensitive to formaldehyde exposure (53). It is possible that both sites being acetylated completely turns off gene expression, while only one leads to intermediate levels of expression, which will be further explored. There were only a few significant changes for *hbsK3Q, hbsK37Q,* and *hbsK86Q* strains and down regulation of *cotD and sspB* at T_4_ were common among them. These mutants were also more susceptible to heat exposure than wildtype after 20 and 30 minutes. Our data suggests that *sspB* and, possibly, *cotD* may contribute to heat resistance. This correlates with previously published findings on reduced resistance of α‾ß‾ SASP mutant spores to dry heat, compared to the wildtype (61). CotD assembles to the inner spore coat and spores that lacked an inner coat were extremely sensitive to lysozyme (62). However, the role of CotD in spore heat resistance is unknown.

There are two known deacetylases (KDACs) in *B. subtilis,* and we previously identified five novel HBsu acetyltransferases (KATs,(57, 63)). In our previous study, we observed a significant decrease in sporulation frequency of the deletion mutant strains Δ*yfmK*, Δ*ydgE* and a double-mutant Δ*yfmK*Δ*ydgE*, whereas these mutations did not affect the resistance properties of the spores (53). These KATs are likely not solely responsible for acetylation at K41, as if there were true, we would expect to see no decrease in sporulation frequency, like observed in the *hbsK41R* strain (53). However, it seems like a sporulation-specific KAT exists, that is activated during early sporulation and advances the cells through the sporulation process by specifically acetylating K41, and possibly other sites. The KATs might have different target sites of HBsu, which was suggested by our previous work (57), but needs to be confirmed. If there were more than one KAT, they could be expressed at different times, which could help the sporulation process continue. It is also possible that they may have additional targets. If there are sporulation-induced KATs, then as the sporulation process advances, a deacetylase would have to be activated to help transition the cells between early and late stages of sporulation. These possibilities are being actively explored in the lab. Together, this data strongly supports the existence of a histone-like code in bacteria and further studies to decipher how bacteria set, erase, and read this code are warranted.

Currently, there are declining options for antibacterial treatment, and these drugs are ineffective against spores, as they are dormant and highly resilient (64, 65, 66). Therefore, continuous research to find new targets and classes of antibiotics with mechanisms of action that could prevent sporulation or promote spore elimination is of interest (67). Disruption of the sporulation-specific gene expression program likely leads to asynchronicity in the intercompartment signaling, which disorganizes the transcriptional program and leads to deficiencies in the final product. We showed that the acetylation state of HBsu is essential for proper gene expression and subsequent resistance characteristics of spores. As a future direction, synthesis of the new small molecule inhibitors that would specifically target HBsu acetylation, either a specific KAT, KDAC, or HBsu itself, could become the foundation for a new class of antibacterial agents that impair sporulation frequency and spore resistance properties by altering the acetylation state of the histone-like proteins.

## Materials and methods

### Bacterial strains, media, and growth conditions

All *B. subtilis* strains are listed in Table 1. The *hbs* point mutations were previously constructed at the native locus, as previously described (55). LB (Luria-Bertani) agar and Schaeffer’s sporulation media (DSM) were prepared as described previously (68). Strains were streaked out on LB plates and incubated overnight at 30°C. The following morning, cells were inoculated into 20 mL of DSM and grown at 37°C with aeration for 6 hours, with growth monitored by Klett colorimetry. 1 mL of cells were harvested after 2, 3, 4, and 6 hours, corresponding to the entry into stationary phase (T_0_), and hours after (T_1_, T_2_, and T_4_ time points), which correlates to stages of sporulation. The samples were centrifuged at 13,000 *x g* for 3 minutes. The resulting cell pellets were processed further either for RT-qPCR, western blot, or microscopy.

### Quantitative real-time polymerase chain reaction (RT-qPCR)

Samples collected at each timepoint were processed with using the FastRNA™ Pro Blue Kit (MP Biomedicals) to extract total RNA as per manufacturer’s instructions, with modifications. Briefly, the cell pellets were resuspended in 1 mL RNApro solution and lysed in tubes containing Lysing Matrix B provided in the kit. The sample tubes were processed using the FastPrep-25 5G Instrument (MP Biomedicals) for three cycles to ensure lysis, 45 sec for the first two, then one 30 sec cycle. The concentration of total RNA was measured using a NanoDrop 2000 spectrophotometer (ThermoFisher Scientific). Average RNA concentrations obtained varied from 800 to 3,500 ng/µL, with an A260:A280 ratio between 1.8-2.1. Next, RNA samples were treated using the Turbo DNA-*free*™ Kit (Invitrogen) according to the manufacturer’s instructions, to remove potential contaminating genomic DNA. Reverse transcription was carried out using the SuperScript™ IV VILO™ kit (Invitrogen) as per manufacturer’s instructions, with the following modifications. The annealing time was extended to 20 minutes at 25°C and the reverse transcription time to 20 minutes at 50°C. The resulting cDNA was purified using the MinElute PCR Purification Kit (Qiagen) according to the manufacturer’s protocol with the following modification. To elute DNA, 12 µL of elution buffer was added to the spin column containing sample, incubated for 5 minutes at 37°C, and then centrifuged at 12,000 *x g* for 2 min. The cDNA samples were diluted 1:5 in nuclease-free water (1X concentration) and the concentrations measured using the NanoDrop 2000. For standard curves, the cDNA was diluted 5-fold six times. 5 μL of 1X, 0.2X, 0.04X, 0.008X, 0.0016X, and 0.00032X concentrations of cDNA for standards and 5 μL of 0.1X for samples, were added to a MicroAmp Optical 96-well plate (Applied Biosystems).12.5 µL of PowerTrack SYBR Green (Applied Biosystems) and 0.25 µL of each primer (10µM) were added to each well, for a 25 µl total volume. All primers used in this study are listed in Table S1. The primers were designed using Primer3 (version 4.1.0), and synthesized by Eton Biosciences (Union, NJ). Plates were run on QuantStudio 7 Pro Real-Time PCR instrument (Applied Biosystems) and analyzed using the relative standard curve method by the built in QuantStudio Design and Analysis software (Applied Biosystems). All reactions were run in technical duplicates, and each experiment was repeated at least three independent times. The *rpoD* transcript was used as an internal control to adjust for differing amounts of input cDNA. All data was normalized to *rpoD* and then the mutants all normalized wildtype (69, 70). A two-tailed T-test was performed to compare relative gene expression of each mutant compared to the wild-type strain, using Microsoft Excel. A p-value of < 0.05 was considered significant. Figures displaying normalized gene expression were created using GraphPad Prism (Version 9.4.0).

### Western Blot

1 mL of cells was collected by centrifugation after 4 and 6 hours of growth in DSM at 37°C. The cell pellets were resuspended in STM buffer (0.71 mM MgCl_2_, 7.14 mM NaCl, 73 mM Sucrose, 35.7 mM Tris, pH8) + 350 µg/mL lysozyme with volumes normalized by Klett, and incubated at 37°C for 5 minutes. Samples were boiled 1X cracking buffer (0.225 M Tris-HCl, pH 6.8, 50% glycerol, 5% sodium dodecyl sulfate [SDS], 0.05% bromophenol blue, 1% β-mercaptoethanol) for 10 minutes and an equal amount of protein loaded on a 15% Tris-Glycine gel. Proteins were transferred to a nitrocellulose membrane using the iBlot 3 Western Blot Transfer System (Invitrogen), according to the manufacturer’s instructions. Membranes were blocked in 5% milk in Tris-buffered saline containing 1% Tween-20 (TBS-T) for 1 hour and then incubated overnight at 4°C with the indicated primary antibodies, at a 1:5,000 dilution for anti-SigF, anti-SigK, or anti-SpoIIAB, and a 1:10,000 dilution for anti-SigG. A 1:5,000 dilution of anti-EFG was included as a loading control. The anti-SigF and anti-SigG antibodies were kindly provided by D. Rudner (Harvard Medical School) and the anti-SigK and anti-SpoIIAB antibodies were kindly provided by N. Bradshaw (Brandeis University). The following day, the membranes were washed four times in TBS-T, then incubated in anti-rabbit secondary antibodies conjugated to horseradish peroxidase (Invitrogen) at a1:5000 dilution for 1 hour at room temperature and washed in TBS-T as before. Bands were visualized using the ECL Prime Western Blotting Detection Reagent (Cytiva) according to the manufacturer’s instructions and imaged using the ChemiDoc™ MP Imaging System (BIO-RAD). All western blots were quantified using Image J (NIH), and all signal intensities were normalized EF-G. To calculate fold changes compared to wildtype, EF-G normalized intensities in the mutant strains were divided by the EF-G normalized intensities in wildtype.

### Sporulation staging using super-resolution microscopy

The cell cultures of BD630 strain were grown to T_0_, T_1_, T_2_, and T_4_ as described before. 5 mL of culture was harvested and centrifuged at 13,000 *x g* for 3 minutes. The supernatant was discarded, and the cell pellets resuspended in 5 mL of phosphate buffered saline (PBS). Cells were washed once with 1 mL of PBS, centrifuged at 13,000 *x g* for 1 minute, and fixed in 1 mL of 4% formaldehyde for 15 min at room temperature. Following incubation, pellets were washed twice with 1mL of PBS, and 200 µL plated on sterile, Lab-Tek II Chambered, #1.5 German cover glass slides (ThermoFisher Scientific), that were previously coated with poly-lysine. Slides were prepared by adding 200 µL of poly-lysine to the slides and incubating them for 20 minutes at room temperature. Slides were then washed with sterile PBS three times for 2 minutes. Membrane permeabilization was performed by adding 200 µL of 2 µg/mL lysozyme for 1 minute, followed by immediate washing with 200 µL of PBS, for a total of three times. Cells on the slides were stained for 5 minutes with 2 µg/ml of Sytox to stain DNA, washed with 200 µL of PBS, for a total of three times, and then stained for 15 minutes with either 10 µg/mL of wheat germ agglutinin conjugated with Alexa Flour 488 (WGA) or 2 µg/mL of Nile Red (Invitrogen). Following staining, slides were washed three times with PBS and imaged on a Nanoimager super-resolution microscope (ONI). The laser power was set at 10-15% for the 488 nm wavelength for Sytox and WGA, and to 10-15% for the 640 nm wavelength for Nile Red. Samples were analyzed using NimOS (version 1.19.4). To determine the ratio of cells in each sporulation stage, at least 200 cells from each time point were counted from three samples of biological replicates.

### Determination of sporulation frequency

The sporulation frequency was performed as previously described(71) with the following modifications. Samples were collected after 2 hours of incubation in DSM (T_0_). The sporulation frequency was calculated as heat-resistant colony-forming units (CFUs) per ml/total viable cells (CFUs/ml). The experiments were carried out at least three independent times. The two-tail Student’s t-test was used to compare the sporulation frequency between samples. A p-value < 0.05 was considered significant.

## Acknowledgements

We thank D. Rudner and N. Bradshaw for kindly providing antibodies.

## References

1. McKenney PT, Driks A, Eichenberger P. The *Bacillus subtilis* endospore: assembly and functions of the multilayered coat. Nat Rev Microbiol. 2013;11(1):33–44.

2. Price EP, Seymour ML, Sarovich DS, Latham J, Wolken SR, Mason J, et al. Molecular epidemiologic investigation of an anthrax outbreak among heroin users, Europe. Emer Infect Dis. 2012;18(8):1307.

3. Tan IS, Ramamurthi KS. Spore formation in *Bacillus subtilis*. Environ Microbiol Rep. 2014;6(3):212–25.

4. Errington J. *Bacillus subtilis* sporulation: regulation of gene expression and control of morphogenesis. Microbiol Rev. 1993;57(1):1–33.

5. Chai Y, He Y, Qin Y, Greenwich J, Balaban S, Darcera MVL, et al. A novel regulation on the developmental checkpoint protein Sda that controls sporulation and biofilm formation in *Bacillus subtilis*. J Bacteriol. 2025, 207 (3), e00210–24.

6. Stragier P, Losick R. Molecular genetics of sporulation in *Bacillus subtilis*. Ann Rev Gen. 1996;30(1):297–341.

7. Fujita M, González-Pastor JE, Losick R. High-and low-threshold genes in the Spo0A regulon of *Bacillus subtilis*. J Bacteriology. 2005;187(4):1357–68.

8. Fujita M, Losick R. Evidence that entry into sporulation in *Bacillus subtilis* is governed by a gradual increase in the level and activity of the master regulator Spo0A. Genes Dev. 2005;19(18):2236–44.

9. Piggot PJ, Hilbert DW. Sporulation of *Bacillus subtilis*. Curr Opin Microbiol. 2004;7(6):579–86.

10. Fimlaid KA, Shen A. Diverse mechanisms regulate sporulation sigma factor activity in the Firmicutes. Curr Opin Microbiol. 2015;24:88–95.

11. Arigoni F, Duncan L, Alper S, Losick R, Stragier P. SpoIIE governs the phosphorylation state of a protein regulating transcription factor sigma F during sporulation in *Bacillus subtilis*. Proc Nat Acad Sci. 1996;93(8):3238–42.

12. Gholamhoseinian A, Piggot PJ. Timing of spoII gene expression relative to septum formation during sporulation of *Bacillus subtilis*. J Bacteriol. 1989;171(10):5747–9.

13. Duncan L, Losick R. SpoIIAB is an anti-sigma factor that binds to and inhibits transcription by regulatory protein sigma F from *Bacillus subtilis*. Proc Nat Acad Sci. 1993;90(6):2325–9.

14. Min K-T, Hilditch CM, Diederich B, Errington J, Yudkin MD. σF, the first compartment-specific transcription factor of *B. subtilis*, is regulated by an anti-σ factor that is also a protein kinase. Cell. 1993;74(4):735–42.

15. Duncan L, Alper S, Arigoni F, Losick R, Stragier P. Activation of cell-specific transcription by a serine phosphatase at the site of asymmetric division. Science. 1995;270(5236):641-4.

16. Levdikov VM, Blagova EV, Rawlings AE, Jameson K, Tunaley J, Hart DJ, et al. Structure of the phosphatase domain of the cell fate determinant SpoIIE from *Bacillus subtilis*. J Molecular Biol. 2012;415(2):343–58.

17. Carniol K, Ben-Yehuda S, King N, Losick R. Genetic dissection of the sporulation protein SpoIIE and its role in asymmetric division in *Bacillus subtilis*. J Bacteriol. 2005;187(10):3511–20.

18. Juan Wu L, Errington J. Identification and characterization of a new prespore-specific regulatory gene, *rsfA*, of *Bacillus subtilis*. J Bacteriol. 2000;182(2):418–24.

19. Ju J, Luo T, Haldenwang WG. *Bacillus subtilis* Pro-sigmaE fusion protein localizes to the forespore septum and fails to be processed when synthesized in the forespore. J Bacteriol. 1997;179(15):4888–93.

20. Imamura D, Zhou R, Feig M, Kroos L. Evidence that the *Bacillus subtilis* SpoIIGA protein is a novel type of signal-transducing aspartic protease. J Biol Chem. 2008;283(22):15287–99.

21. McBride S, Haldenwang WG. Sporulation phenotype of a *Bacillus subtilis* mutant expressing an unprocessable but active sigmaE transcription factor. J Bacteriol. 2004;186(7):1999–2005.

22. Dworkin J, Losick R. Developmental commitment in a bacterium. Cell. 2005;121(3):401–9.

23. Narula J, Devi SN, Fujita M, Igoshin OA. Ultrasensitivity of the *Bacillus subtilis* sporulation decision. Proc Natl Acad Sci U S A. 2012;109(50):E3513–22.

24. Schumacher MA, Lee J, Zeng W. Molecular insights into DNA binding and anchoring by the Bacillus subtilis sporulation kinetochore-like RacA protein. Nucleic Acids Res. 2016;44(11):5438–49.

25. Doan T, Marquis KA, Rudner DZ. Subcellular localization of a sporulation membrane protein is achieved through a network of interactions along and across the septum. Mol Microbiol. 2005;55(6):1767–81.

26. Gutierrez J, Smith R, Pogliano K. SpoIID-mediated peptidoglycan degradation is required throughout engulfment during *Bacillus subtilis* sporulation. J Bacteriol. 2010;192(12):3174–86.

27. Sun DX, Cabrera-Martinez RM, Setlow P. Control of transcription of the *Bacillus subtilis spoIIIG* gene, which codes for the forespore-specific transcription factor sigma G. J Bacteriol. 1991;173(9):2977–84.

28. Driks A, Eichenberger P. The sporecoat. Microbiol Spectr. 2016;4(2).

29. McKenney PT, Eichenberger P. Dynamics of spore coat morphogenesis in *Bacillus subtilis*. Mol Microbiol. 2012;83(2):245–60.

30. Chary VK, Meloni M, Hilbert DW, Piggot PJ. Control of the expression and compartmentalization of (sigma)G activity during sporulation of *Bacillus subtilis* by regulators of (sigma)F and (sigma)E. J Bacteriol. 2005;187(19):6832–40.

31. Serrano M, Neves A, Soares CM, Moran CP, Jr., Henriques AO. Role of the anti-sigma factor SpoIIAB in regulation of sigmaG during *Bacillus subtilis* sporulation. J Bacteriol. 2004;186(12):4000–13.

32. Hilbert DW, Piggot PJ. Compartmentalization of gene expression during *Bacillus subtilis* spore formation. Microbiol Mol Biol Rev. 2004;68(2):234–62.

33. Moeller R, Setlow P, Reitz G, Nicholson WL. Roles of small, acid-soluble spore proteins and core water content in survival of *Bacillus subtilis* spores exposed to environmental solar UV radiation. Appl Environ Microbiol. 2009;75(16):5202–8.

34. Nerber HN, Sorg JA. The small acid-soluble proteins of spore-forming organisms: similarities and differences in function. Anaerobe. 2024;87:102844.

35. Setlow B, Setlow P. Binding of small, acid-soluble spore proteins to DNA plays a significant role in the resistance of *Bacillus subtilis* spores to hydrogen peroxide. Appl Environ Microbiol. 1993;59(10):3418–23.

36. Wang ST, Setlow B, Conlon EM, Lyon JL, Imamura D, Sato T, et al. The forespore line of gene expression in *Bacillus subtilis*. J Mol Biol. 2006;358(1):16–37.

37. Haldenwang WG. The sigma factors of *Bacillus subtilis*. Microbiol Rev. 1995;59(1):1–30.

38. Meador-Parton J, Popham DL. Structural analysis of *Bacillus subtilis* spore peptidoglycan during sporulation. J Bacteriol. 2000;182(16):4491–9.

39. Higgins D, Dworkin J. Recent progress in *Bacillus subtilis* sporulation. FEMS Microbiol Rev. 2012;36(1):131–48.

40. Ross MA, Setlow P. The *Bacillus subtilis* HBsu protein modifies the effects of alpha/beta-type, small acid-soluble spore proteins on DNA. J Bacteriol. 2000;182(7):1942–8.

41. Cortesao M, Fuchs FM, Commichau FM, Eichenberger P, Schuerger AC, Nicholson WL, et al. *Bacillus subtilis* spore resistance to simulated mars surface conditions. Front Microbiol. 2019;10:333.

42. Setlow P. Spore resistance properties. Microbiol Spectr. 2014;2(5).

43. Magge A, Granger AC, Wahome PG, Setlow B, Vepachedu VR, Loshon CA, et al. Role of dipicolinic acid in the germination, stability, and viability of spores of *Bacillus subtilis*. J Bacteriol. 2008;190(14):4798–807.

44. Mason JM, Setlow P. Essential role of small, acid-soluble spore proteins in resistance of *Bacillus subtilis* spores to UV light. J Bacteriol. 1986;167(1):174–8.

45. Moeller R, Setlow P, Horneck G, Berger T, Reitz G, Rettberg P, et al. Roles of the major, small, acid-soluble spore proteins and spore-specific and universal DNA repair mechanisms in resistance of *Bacillus subtilis* spores to ionizing radiation from X rays and high-energy charged-particle bombardment. J Bacteriol. 2008;190(3):1134–40.

46. Alonso JC, Gutierrez C, Rojo F. The role of the chromatin-associated protein HBsu in beta-mediated DNA recombination is to facilitate the joining of distant recombination sites. Mol Microbiol. 1995;18(3):471–8.

47. Fernandez S, Rojo F, Alonso JC. The *Bacillus subtilis* chromatin-associated protein HBsu is involved in DNA repair and recombination. Mol Microbiol. 1997;23(6):1169–79.

48. Alonso JC, Weise F, Rojo F. The B*acillus subtilis* histone-like protein HBsu is required for DNA resolution and DNA inversion mediated by the beta-recombinase of plasmid Psm19035. J Biol Chem. 1995;270(7):2938–45.

49. Karaboja X, Wang X. HBsu Is required for the initiation of DNA replication in *Bacillus subtilis*. J Bacteriol. 2022;204(8):e0011922.

50. Köhler P, Marahiel MA. Association of the histone-like protein HBsu with the nucleoid of *Bacillus subtilis*. J Bacteriol. 1997;179(6):2060–4.

51. Christensen DG, Baumgartner JT, Xie X, Jew KM, Basisty N, Schilling B, et al. Mechanisms, detection, and relevance of protein acetylation in prokaryotes. mBio. 2019;10(2).

52. Luu J, Carabetta VJ. Contribution of N(epsilon)-lysine acetylation towards regulation of bacterial pathogenesis. mSystems. 2021;6(4):e0042221.

53. Luu J, Mott CM, Schreiber OR, Giovinco HM, Betchen M, Carabetta VJ. N(epsilon)-Lysine acetylation of the histone-like protein HBsu regulates the process of sporulation and affects the resistance properties of *Bacillus subtilis* spores. Front Microbiol. 2021;12:782815.

54. Carabetta VJ, Greco TM, Tanner AW, Cristea IM, Dubnau D. Temporal regulation of the *Bacillus subtilis* acetylome and evidence for a role of MreB acetylation in cell wall growth. mSystems. 2016;1(3).

55. Carabetta VJ, Greco TM, Cristea IM, Dubnau D. YfmK is an N(epsilon)-lysine acetyltransferase that directly acetylates the histone-like protein HBsu in *Bacillus subtilis*. Proc Natl Acad Sci. 2019;116(9):3752–7.

56. Wang ST, Setlow B, Conlon EM, Lyon JL, Imamura D, Sato T, et al. The forespore line of gene expression in *Bacillus subtilis*. J Mol Biol. 2006;358(1):16–37.

57. Carabetta VJ, Greco TM, Cristea IM, Dubnau D. YfmK is an Nε-lysine acetyltransferase that directly acetylates the histone-like protein HBsu in *Bacillus subtilis*. Proc Natl Acad Sci. 2019;116(9):3752–7.

58. Micka B, Marahiel MA. The DNA-binding protein HBsu is essential for normal growth and development in *Bacillus subtilis*. Biochimie. 1992;74(7-8):641–50.

59. Klein W, Marahiel MA. Structure-function relationship and regulation of two *Bacillus subtilis* DNA-binding proteins, HBsu and AbrB. J Mol Microbiol Biotechnol. 2002;4(3):323–9.

60. Carr RA, Tucker T, Newman PM, Jadalla L, Jaludi K, Reid BE, et al. N(epsilon)-lysine acetylation of the histone-like protein HBsu influences antibiotic survival and persistence in *Bacillus subtilis*. Front Microbiol. 2024;15:1356733.

61. Setlow B, Setlow P. Small, acid-soluble proteins bound to DNA protect *Bacillus subtilis* spores from killing by dry heat. Appl Environ Microbiol. 1995;61(7):2787–90.

62. Riesenman PJ, Nicholson WL. Role of the spore coat layers in *Bacillus subtilis* spore resistance to hydrogen peroxide, artificial UV-C, UV-B, and solar UV radiation. Appl Environ Microbiol. 2000;66(2):620–6.

63. Gardner JG, Escalante-Semerena JC. In *Bacillus subtilis*, the sirtuin protein deacetylase, encoded by the *srtN* gene (formerly *yhdZ)*, and functions encoded by the *acuABC* genes control the activity of acetyl coenzyme A synthetase. J Bacteriol. 2009;191(6):1749–55.

64. Butler MS, Henderson IR, Capon RJ, Blaskovich MAT. Antibiotics in the clinical pipeline as of December 2022. J Antibiot (Tokyo). 2023;76(8):431–73.

65. Louie A, VanScoy BD, Brown DL, Kulawy RW, Heine HS, Drusano GL. Impact of spores on the comparative efficacies of five antibiotics for treatment of *Bacillus anthracis* in an in vitro hollow fiber pharmacodynamic model. Antimicrob Agents Chemother. 2012;56(3):1229–39.

66. Romero-Rodriguez A, Ruiz-Villafan B, Martinez-de la Pena CF, Sanchez S. Targeting the impossible: A review of new strategies against endospores. Antibiotics. 2023;12(2).

67. Andryukov BG, Karpenko AA, Lyapun IN. Learning from nature: Bacterial spores as a target for current technologies in medicine (Review). Sovrem Tekhnologii Med. 2021;12(3):105–22.

68. Albano M, Hahn J, Dubnau D. Expression of competence genes in *Bacillus subtilis*. J Bacteriol. 1987;169(7):3110–7.

69. Biosystems A. Guide to performing relative quantitation of gene expression using real-time quantitative PCR. Applied biosystems. 2004.

70. Larionov A, Krause A, Miller W. A standard curve based method for relative real time PCR data processing. BMC Bioinformatics. 2005;6:1–16.

71. Carabetta VJ, Tanner AW, Greco TM, Defrancesco M, Cristea IM, Dubnau D. A complex of YlbF, YmcA and YaaT regulates sporulation, competence and biofilm formation by accelerating the phosphorylation of Spo0A. Mol Microbiol. 2013;88(2):283–300.

