## Supplemental figures S1-S7, tables S1-S3 for "The acetylation of the histone-like protein HBsu at specific sites alters gene expression during sporulation in *Bacillus subtilis*"

**Figure S1**. Interplay of sporulation-specific σ and transcription factors in *B. subtilis*

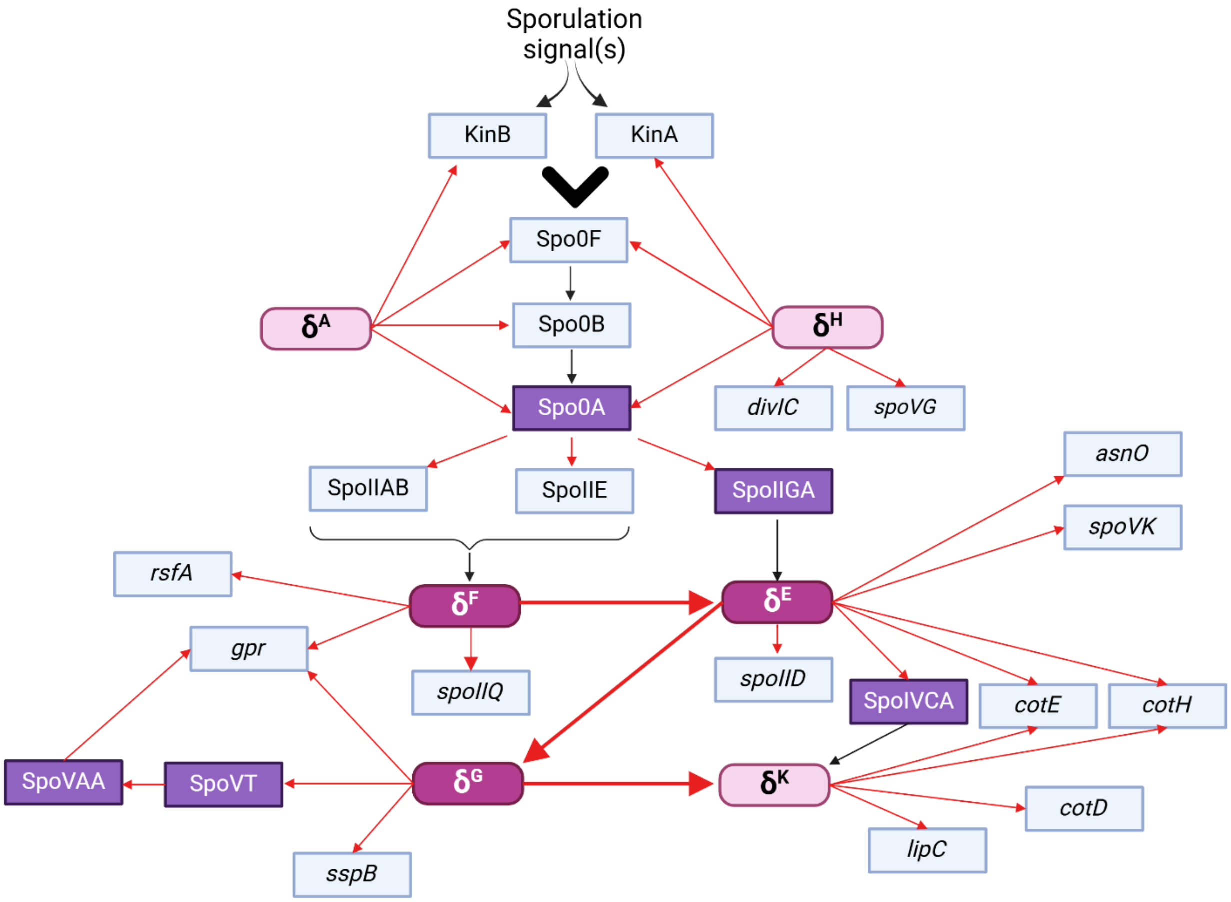

The network displays the interactions among the genes, and their products, that were analyzed in this study. The sigma factors that were directly assayed are dark pink, and the others (light pink) were assayed for activity by monitoring expression of their regulon members. Transcription factors are indicated in purple. Nodes connected by red arrows represent a transcriptional relationship, and those connected by black arrows represent post-translational regulation. The large arrows represent sigma factor cross-compartment signaling. Activation of Spo0A occurs through the phosphorelay, and expression of these genes are under the control of $\sigma^A$ *and* $\sigma^H$ regulons (1, 2). The $\sigma^H$ regulon also includes *divIC,* which encodes a polar division protein and *spoVG,* which controls asymmetric septation and later cortex formation (3, 4, 5). To activate sporulation, Spo0F is phosphorylated by one of several histidine kinases, including KinA and KinB, which in turn leads to the transfer of phosphate to Spo0B, and ultimately Spo0A (1, 2). Spo0A~P activates transcription of the genes responsible for entry into sporulation, including *spoIIE*, *spoIIA* and *spoIIG* operons (2). SpoIIE and SpoIIAB are involved in the $\sigma^F$ -activation cascade (6, 7, 8, 9, 10). SpoIIGA catalyses the cleavage of 27 amino acids from the pro- $\sigma^E$ amino terminus and, thereby, $\sigma^E$ activation (11, 12). The $\sigma^F$ regulon include *rsf* (13, 14), *spoIIQ* and *gpr* (14). The $\sigma^E$ regulon includes *asnO* (15), *spoIID, spoVK* (16), *spoIVCA, cotE* and *cotH* (17, 18, 19). Overall, $\sigma^F$ and $\sigma^E$ control subsequent sporulation steps, like engulfment, which depends on SpoIID and SpoIIQ (20, 21), and activation of the late sporulation-specific sigma factors $\sigma^G$ and $\sigma^K$. $\sigma^G$ controls the transcription of gpr, *sspB,* and *spoVT*, which itself is a transcription factor that regulates the expression *spoVAA* (22, 23). SpoIVCA excises the *skin* element, to lead to $\sigma^K$ activation (24). $\sigma^K$ regulates the expression of *cotE, cotH, cotD* (25, 26), and *lipC* (27).

**Figure S2.** Staging of wild-type *B. subtilis* cells in DSM.

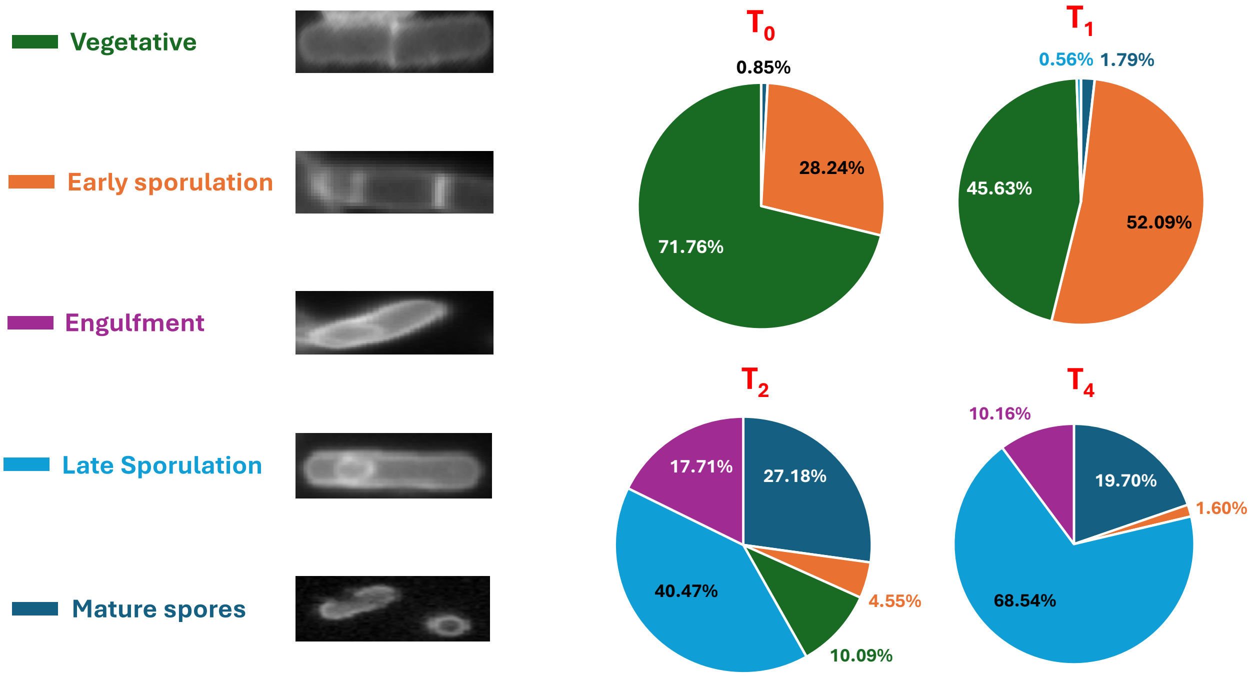

Wild-type cells were grown in DSM, with time points taken from $T_0$-$T_4$. Cells were collected, fixed, and seeded on chambered, cover glass slides coated with poly-lysine. Cells were stained with wheat germ agglutinin conjugated with Alexa Flour 488 or Nile red, and analyzed by super-resolution microscopy, as described in Materials and Methods. The staging was determined using different biological replicates, analyzing at least 250 cells in each sample. At $T_0$ most cells are in vegetative growth (71.76%), whereas only 28.24% of the cells transitioned to sporulation. At $T_1$ more than a half of the cell population (52.9%) transitioned into sporulation, and 45.63% were vegetative. At $T_2$ we observed cells later in the sporulation process, including engulfment (17.71%) and into late sporulation (40.47%). In addition, some mature spores were observed. At $T_4$ all cells had entered sporulation and were at various stages. Most cells (88.24%) were observed in late sporulation, including ~20% spores. This confirms that in the cells progress through the sporulation process as expected in our laboratory media, and we are examining the appropriate time points.

**Figure S3.** SpoIIAB levels in wild-type and *hbsK41Q* strains.

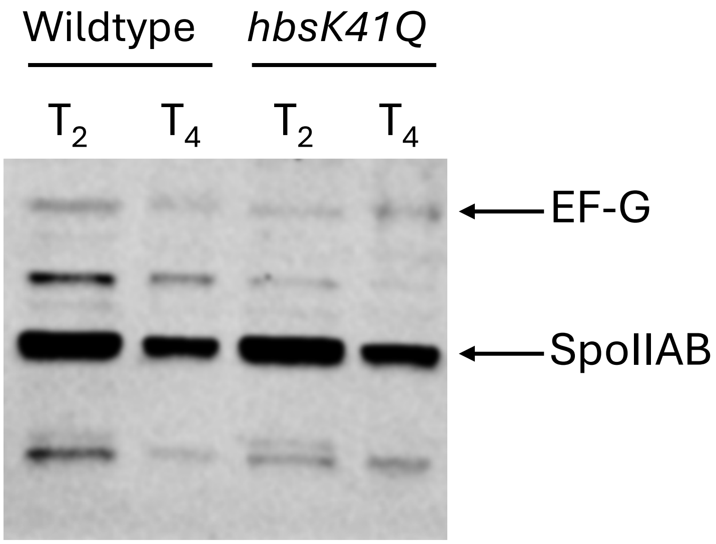

Wild-type and *hbsK41Q* cells were grown to $T_2$ and $T_4$ in sporulation media and lysates prepared as described in Materials and Methods. Equal amounts of protein were loaded and were probed with anti-SpoIIAB and anti-EFG antibodies. EFG was included as a loading control. Non-labeled bands represent cross-reacting bands. All western blots were repeated three independent times, and a representative blot is shown.

**Figure S4.** Unmodified versions of Western blot in figure 5 at different exposures.

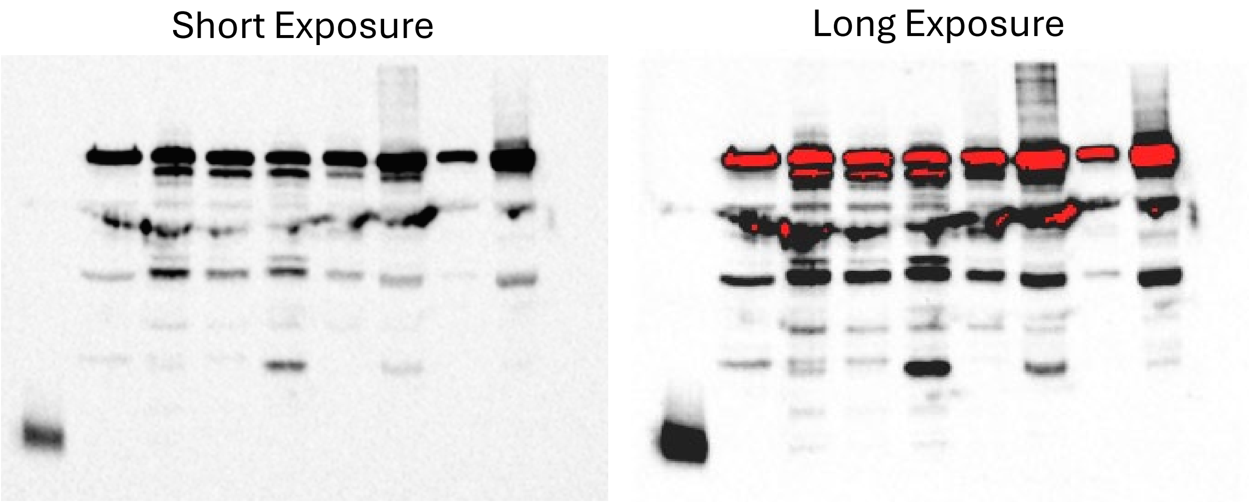

Full, uncropped versions of the western blot in Fig. 5. The short exposure was for 15 sec, which was necessary to visualize the EF-G signal. The longer exposure was for 60 sec to visualize the $\sigma^G$ band. The instrument false-colored bands in red that were saturated and out of the linear range of detection. The EF-G band is saturated at this exposure, necessitating analysis at different exposure times for each signal.

**Figure S5.** Sporulation frequences at $T_0$ under sporulation conditions.

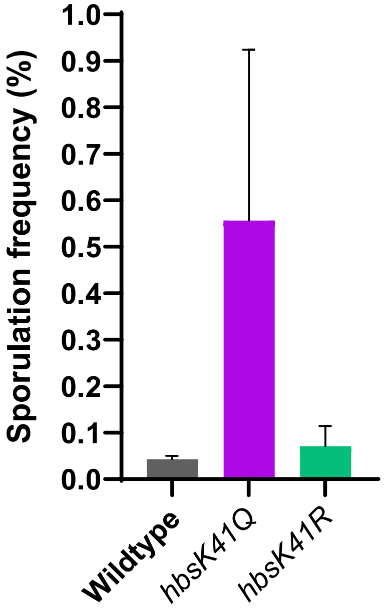

Wild-type, *hbsK41Q,* and *hbsK41R* cells were grown for 2 hours in DSM media to $T_0$. The displayed values are the mean of three biological replicates, with standard deviation shown. For the wild-type and *hbsK41R* strains, there was little sporulation, but detectable levels of sporulation in the *hbsK41Q* strain. None of the differences were statistically significant.

**Figure S6.** Gene expression changes in *hbsK37Q* strain.

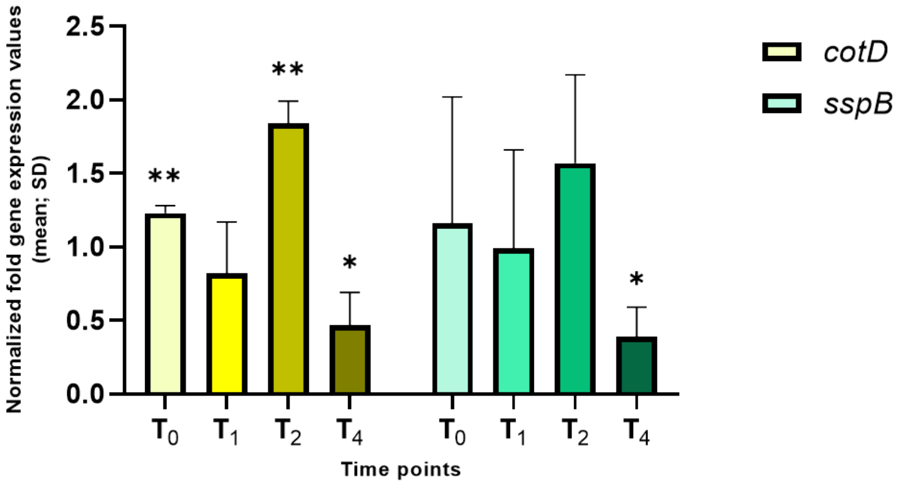

Wildtype and the *hbsK37Q* mutant were grown in DSM for 6 hours and analyzed at time points $T_0$-$T_4$. Values displayed are the mean expression of technical duplicates from at least three biological replicates with the standard deviation (SD). The reduction in expression of *cotD* (coat protein) and *sspB* (ß-type small acid-soluble protein) at $T_4$ might explain the higher susceptibility to heat. *$p<0.05$, ** $p<0.01$

**Figure S7.** Altered gene expression in *hbsK86Q* mutants.

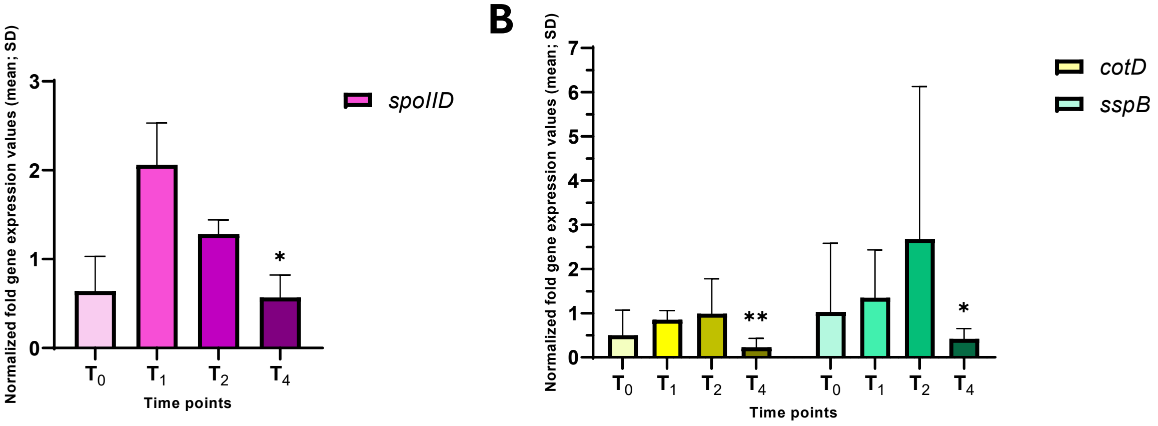

Wild-type and *hbsK86Q* cells were grown in DSM for 6 hours and analyzed at time points $T_0$-$T_4$. Values displayed are the mean expression of technical duplicates from at least three biological replicates with the standard deviation (SD). At $T_4$ there was a > 2-fold reduction of *cotD* and *sspB*, and a reduction of *spoIID* expression. Reduction of these three genes may contribute to the decreased resistance properties of mutant spores. *p<0.05, ** p<0.01

**Table S1.** Primers used in this study

| Primer name | Sequence (5' → 3') |
|---|---|
| *5-cotD-RT* | TGGCGCCAATTGTCCATCCT |
| *3-cotD-RT* | ACGTGCTGAAAATGCTGGTGG |
| *5-spoIID-RT* | CAGTCAGCACGAAACCAGCA |
| *3-spoIID-RT* | AGGCGACGACTCCAATCACA |
| *5-sspB-RT* | ACCTTGGAGCGGACACAACT |
| *3-sspB-RT* | TGAACTCTGCCGCCCATTTG |
| *5-spoVG-RT* | CGATTGCATCCATCACGCTG |
| *3-spoVG-RT* | ATCAGGGGTGCGTTTACTCG |
| *5-spo0F-RT* | TGGACATGAAAATTCCCGGCA |
| *3-spo0F-RT* | AGAGCGCCCAATTCCTTCGA |
| *5-spo0A-RT* | TCCGCCATGCAATTGAAGTGG |
| *3-spo0A-RT* | ACCTCAGCTTATCCGCAACCA |
| *5-kinB-RT* | ACCTGCGATTCTTGTGCCAA |
| *3-kinB-RT* | AACGGGAATCATCTGAAGGCC |
| *5-spoIIGA-RT* | GGAACAGCCGAAACGATGATCA |
| *3-spoIIGA-RT* | TTGCTGACCAACTCCCCTGT |
| *5-spo0B-RT* | TCATCTGCTTGGCCATTCCC |
| *3-spo0B-RT* | GCTTTGATTCGTGCTTTGCGT |
| *5 KinA RT* | TGAGCGGACAGAACGGGAAA |
| *3 KinA RT* | CGGGCTTGCAGGTTCGATTT |
| *5 spoIIE RT* | TTCCGGGAACTGTCGAGCAT |
| *3 spoIIE RT* | GCTTTCAGACAGCGCGTGAA |
| *5 spoIIAB RT* | AGCTGGACCCGACAATGGAT |
| *3 spoIIAB RT* | ACGACATGATCTTCCAGCGTCA |
| *5 sigF RT* | GCGTCTTGTTTGGTCTGTCGT |
| *3 sigF RT* | TCGGCACTGCATACGTTGAA |
| *5 rsfA RT* | TCCGTGCTTGAGAGTGATGC |
| *3 rsfA RT* | GGATCACCGTTCGCCACAAT |
| *5 sigG RT* | AGAGGGCATGAGAAGGCTGA |
| *3 sigG RT* | TGGACACCTGCGCTTGAGAA |
| *5 gpr RT* | TGACACCTGATGCGCTTGGA |
| *3 gpr RT* | TGCAAAAGCACTGACAGGCC |
| *5 spoIIQ RT* | TAGTCAGTGCGGCCGTCATT |
| *3 spoIIQ RT* | ACTGCATCGTCGTTGTTGTCA |
| *5 spoVK RT* | ACACCGCCCAAAAGACGAGA |
| *3 spoVK RT* | AAGTCCTTTTCGCCGCCTCT |
| *5 asnO RT* | TGTTGTTTGCCGCGAGAGAC |
| *3 asnO RT* | TATCAGGGTGCGCGAGGATT |
| *5 spoIVCA RT* | TCGACCGAGGAACAAGCGAT |

| | |
|---|---|
| *3 spoIVCA RT* | GCGATTCAAAGCCGGACGTT |
| *5 divIC RT* | CCAGGGAACGAACGATAACTGA |
| *3 divIC RT* | ACTAGGGCGCCGAATACAGT |
| *5 spoVAA RT* | TGCCGCTTTATCAGGTGAGC |
| *3 spoVAA RT* | ACAATGGTTTCTGCTCCGCC |
| *5 spoVT RT* | TCCGTGCTTGAGAGTGATGC |
| *3 spoVT RT* | GGATCACCGTTCGCCACAAT |
| *5 cotE RT* | AACCGAGCAGCATTTTGGGT |
| *3 cotE RT* | TGTCTTTGTGTTGTCCGCGT |
| *5 cotH RT* | TTTCTCGGAGCTAGGGACACTG |
| *3 cotH RT* | GCCAGCTTCCTTTTCGCCAA |
| *5 lipC RT* | AAATCCTTCCCCGTACGCCA |
| *3 lipC RT* | CCGCGATTTTGACATGGGGT |
| *5 rpoD RT* | AAAACGGTATGTCGGACGCG |
| *3 rpoD RT* | ATCGCCTGTCTGATCCACCA |
| *5 rpoA RT* | TGCTCAAAGAGGACGTGGGT |
| *3 rpoA RT* | TGCAACTTGGCCTACACGAG |
| *5 rrnA-16S RT* | TGGTTGTCGTCAGCTCGTGT |
| *3 rrnA-16S RT* | TTGTCACCGGCAGTCACCTT |

**Table S2.** Primer efficiency for selected housekeeping genes.

| Gene | Efficiency | Curve | $R^2$ |
|---|---|---|---|
| ***rpoA*** | **79.57%** | -3.94x + 19.8 | 0.9991 |
| ***rpoD*** | **96.58%** | -3.53x + 23.7 | 0.9962 |
| ***rrnA-16S*** | **81.68%** | -3.85x + 12.3 | 0.9898 |
| ***rrnA-5S*** | **54.01%** | -5.33x + 19.5 | 0.9874 |

### **Table S3.** List of selected genes

| Gene | Description* | Function* |
|---|---|---|
| *spo0A* | Sporulation transcription factor Spo0A, a phosphorelay regulator, phosphorylated in response to complex YlbF/YmcA/YaaT. | Initiation of sporulation |
| *spo0B* | Sporulation initiation phosphotransferase Spo0B | Initiation of sporulation |
| *spo0F* | Phosphotransferase, initiation of sporulation phosphorelay | Initiation of sporulation |
| *kinA* | Sporulation-specific, ATP-dependent protein histidine kinase, phosphorylates Spo0F | Initiation of sporulation |
| *kinB* | Sporulation sensor histidine kinase KinB, phosphorylates Spo0F | Initiation of sporulation |
| *spoIIQ* | Forespore protein, part of the transmembrane channel linking the mother cell and the forespore, stage II sporulation protein SpoIIQ | Forespore encasement by the spore coat |
| *spoIIE* | SpoIIAA-phosphate serine phosphatase, stage II sporulation protein E | Control of $\sigma^F$ activity, required for formation of the asymmetric septum |
| *sigF* | RNA polymerase sporulation-specific sigma factor (sigma-F) | transcription of sporulation genes (early forespore) |
| *spoIIAB* | Anti-sigma factor (antagonist of $\sigma^F$) and serine kinase | control of $\sigma^F$ activity; phosphorylation and inactivation of SpoIIAA |
| *rsfA* | Prespore-specific transcription regulator RsfA of $\sigma^F$ activity | Control of expression of $\sigma^F$-dependent genes |
| *sigG* | RNA polymerase sporulation-specific sigma factor (sigma-G) | Transcription of sporulation genes (late forespore) |
| *spoIIGA* | Protease processing pro- $\sigma^E$ | Maturation of $\sigma^E$ |
| *spoIID* | Lytic transglycosylase | Dissolution of the septal cell wall |
| *divIC* | Cell-division initiation protein | Septum formation |
| *spoIVCA* | Site-specific DNA recombinase required for creating the *sigK* gene | Excision of the Skin element, creation of the *sigK* gene |
| *asnO* | Asparagine synthetase (sporulation related) | Biosynthesis of asparagine |
| *cotE* | Morphogenic spore protein, outer spore coat protein CotE | Assembly of the outer spore coat |
| *cotH* | Spore coat protein kinase | Protection of CotU and CotC in the mother cell |
| *cotD* | Spore coat protein (inner) | Resistance of the spore |
| *spoVAA* | Stage V sporulation protein AA | Spore maturation |
| *spoVT* | Transcriptional regulator of sporulation / germination, stage V sporulation protein T | regulation of forespore gene expression |
| *spoVK* | Mother cell sporulation ATPase, stage V sporulation protein K | Spore maturation |
| *spoVG* | RNA-binding regulatory protein, affects asymmetric septation and cortex formation | Cell division, control of sporulation initiation, cortex formation |
| *sspB* | Small acid-soluble spore protein (major beta-type SASP) | Protection of spore DNA |
| *lipC* | Spore coat phospholipase B | Spore germination |
| *gpr* | Spore germination protease, GPR endopeptidase | Degradation of SASPs |

*Descriptions and functions are from *Subtiwiki* (Elfmann C, Dumann V, van den Berg T, Stülke J. A new framework for SubtiWiki, the database for the model organism *Bacillus subtilis*. Nucleic Acids Res. 2025 Jan 6;53(D1):D864-D870. doi: 10.1093/nar/gkae957).

### Supplemental References

1. Levin PA, Losick R. Transcription factor Spo0A switches the localization of the cell division protein FtsZ from a medial to a bipolar pattern in *Bacillus subtilis*. Genes Dev. 1996;10(4):478-88.
2. Stragier P, Losick R. Molecular genetics of sporulation in *Bacillus subtilis*. Ann Rev Gen. 1996;30(1):297-341.
3. Eijlander RT, Holsappel S, de Jong A, Ghosh A, Christie G, Kuipers OP. SpoVT: From fine-tuning regulator in *Bacillus subtilis* to essential sporulation protein in *Bacillus cereus*. Front Microbiol. 2016;7:1607.
4. Katis VL, Harry EJ, Wake RG. The *Bacillus subtilis* division protein DivIC is a highly abundant membrane-bound protein that localizes to the division site. Mol Microbiol. 1997;26(5):1047-55.
5. Ramirez-Peralta A, Stewart KA, Thomas SK, Setlow B, Chen Z, Li YQ, et al. Effects of the SpoVT regulatory protein on the germination and germination protein levels of spores of *Bacillus subtilis*. J Bacteriol. 2012;194(13):3417-25.
6. Carniol K, Ben-Yehuda S, King N, Losick R. Genetic dissection of the sporulation protein SpoIIE and its role in asymmetric division in *Bacillus subtilis*. J Bacteriol. 2005;187(10):3511-20.
7. Duncan L, Alper S, Losick R. SpoIIAA governs the release of the cell-type specific transcription factor sigma F from its anti-sigma factor SpoIIAB. J Mol Biol. 1996;260(2):147-64.
8. Duncan L, Losick R. SpoIIAB is an anti-sigma factor that binds to and inhibits transcription by regulatory protein sigma F from *Bacillus subtilis*. Proc Natl Acad Sci U S A. 1993;90(6):2325-9.
9. Garsin DA, Duncan L, Paskowitz DM, Losick R. The kinase activity of the antisigma factor SpoIIAB is required for activation as well as inhibition of transcription factor sigmaF during sporulation in *Bacillus subtilis*. J Mol Biol. 1998;284(3):569-78.
10. Kellner EM, Decatur A, Moran CP, Jr. Two-stage regulation of an anti-sigma factor determines developmental fate during bacterial endospore formation. Mol Microbiol. 1996;21(5):913-24.
11. Imamura D, Zhou R, Feig M, Kroos L. Evidence that the *Bacillus subtilis* SpoIIGA protein is a novel type of signal-transducing aspartic protease. J Biol Chem. 2008;283(22):15287-99.
12. Ju J, Luo T, Haldenwang WG. *Bacillus subtilis* Pro-sigmaE fusion protein localizes to the forespore septum and fails to be processed when synthesized in the forespore. J Bacteriol. 1997;179(15):4888-93.
13. Wu LJ, Errington J. Identification and characterization of a new prespore-specific regulatory gene, rsfA, of *Bacillus subtilis*. J Bacteriol. 2000;182(2):418-24.
14. Wang ST, Setlow B, Conlon EM, Lyon JL, Imamura D, Sato T, et al. The forespore line of gene expression in *Bacillus subtilis*. J Mol Biol. 2006;358(1):16-37.
15. Yoshida K, Fujita Y, Ehrlich SD. Three asparagine synthetase genes of *Bacillus subtilis*. J Bacteriol. 1999;181(19):6081-91.
16. Delerue T, Chareyre S, Anantharaman V, Gilmore MC, Popham DL, Cava F, et al. Bacterial cell surface nanoenvironment requires a specialized chaperone to activate a peptidoglycan biosynthetic enzyme. bioRxiv. 2023.
17. Bauer T, Little S, Stover AG, Driks A. Functional regions of the *Bacillus subtilis* spore coat morphogenetic protein CotE. J Bacteriol. 1999;181(22):7043-51.
18. Giglio R, Fani R, Isticato R, De Felice M, Ricca E, Baccigalupi L. Organization and evolution of the cotG and cotH genes of *Bacillus* subtilis. J Bacteriol. 2011;193(23):6664-73.
19. Zilhao R, Naclerio G, Henriques AO, Baccigalupi L, Moran CP, Jr., Ricca E. Assembly requirements and role of CotH during spore coat formation in *Bacillus subtilis*. J Bacteriol. 1999;181(8):2631-3.

20. Campo N, Marquis KA, Rudner DZ. SpoIIQ anchors membrane proteins on both sides of the sporulation septum in *Bacillus subtilis*. J Biol Chem. 2008;283(8):4975-82.
21. Nocadello S, Minasov G, Shuvalova LS, Dubrovska I, Sabini E, Anderson WF. Crystal structures of the SpoIID lytic transglycosylases essential for bacterial sporulation. J Biol Chem. 2016;291(29):14915-26.
22. Magge A, Granger AC, Wahome PG, Setlow B, Vepachedu VR, Loshon CA, et al. Role of dipicolinic acid in the germination, stability, and viability of spores of *Bacillus subtilis*. J Bacteriol. 2008;190(14):4798-807.
23. Setlow B, Atluri S, Kitchel R, Koziol-Dube K, Setlow P. Role of dipicolinic acid in resistance and stability of spores of *Bacillus subtilis* with or without DNA-protective alpha/beta-type small acid-soluble proteins. J Bacteriol. 2006;188(11):3740-7.
24. Popham DL, Stragier P. Binding of the *Bacillus subtilis* spoIVCA product to the recombination sites of the element interrupting the sigma K-encoding gene. Proc Natl Acad Sci U S A. 1992;89(13):5991-5.
25. Driks A, Eichenberger P. The spore coat. Microbiol Spectr. 2016;4(2).
26. McKenney PT, Eichenberger P. Dynamics of spore coat morphogenesis in *Bacillus subtilis*. Mol Microbiol. 2012;83(2):245-60.
27. Masayama A, Kuwana R, Takamatsu H, Hemmi H, Yoshimura T, Watabe K, et al. A novel lipolytic enzyme, YcsK (LipC), located in the spore coat of *Bacillus subtilis*, is involved in spore germination. J Bacteriol. 2007;189(6):2369-75.
